# Genotypic variation in soil penetration by maize roots is negatively related to ethylene-induced thickening

**DOI:** 10.1101/2021.01.15.426842

**Authors:** Dorien J. Vanhees, Hannah M. Schneider, Kenneth W. Loades, A. Glyn Bengough, Malcolm J. Bennett, Bipin K. Pandey, Kathleen M. Brown, Sacha J. Mooney, Jonathan P. Lynch

## Abstract

Radial expansion is a classic response of roots to mechanical impedance that has generally been assumed to aid penetration. We analysed the response of maize nodal roots to impedance to test the hypothesis that radial expansion is not related to the ability of roots to cross a compacted soil layer. Genotypes varied in their ability to cross the compacted layer, and those with a steeper approach to the compacted layer or less radial expansion in the compacted layer were more likely to cross the layer and achieve greater depth. Root radial expansion was due to cortical cell size expansion, while cortical cell file number remained constant. Genotypes and nodal root classes that exhibited radial expansion upon encountering the compacted soil layer also thickened in response to exogenous ethylene in hydroponic culture, i.e. radial expansion in response to ethylene was correlated with the thickening response to impedance in soil. We propose that ethylene insensitive roots, i.e. those that do not thicken and are able to overcome impedance, have a competitive advantage under mechanically impeded conditions as they can maintain their elongation rates. We suggest that prolonged exposure to ethylene could function as a stop signal for axial root growth.

## Introduction

Roots interact dynamically with the highly heterogeneous soil environment and commonly need to withstand abiotic and biotic stresses in order to acquire water and nutrients. One major constraint to root growth and function is mechanical impedance, or the physical resistance to root penetration imposed by soil (Bennie, 1996; Whalley *et al*., 2005). An example of localised mechanically impeding conditions that roots encounter is the presence of harder soil clods or aggregates (Konôpka *et al*., 2009, 2008). Another example is plough pans created by tillage which are spatially abrupt. Roots unable to penetrate through harder soil strata run the risk of being confined to the upper, less dense soil domains while roots adapted to impeded conditions are able to penetrate through harder layers and would be able to maintain normal plant growth (Barraclough and Weir, 1988; Ehlers *et al*., 1983; Pfeifer *et al*., 2014). Soil structure itself can facilitate root exploration but could also hinder root growth. Biopores formed by pre-existing roots can be used to bypass harder soil domains (Athmann, 2019; Ehlers *et al*., 1983; Han *et al*., 2015; Valentine *et al*., 2012; Whitmore and Whalley, 2009). However, roots can become confined in soil pores restricting soil exploration of the bulk soil (Pankhurst *et al*., 2002; White and Kirkegaard, 2010). As a localised denser region of soil surrounds a root (Helliwell *et al*., 2019), a pore formed by previous roots might constrict subsequent roots due to greater impedance in the pore wall. In order to further explore bulk soil a root must therefore overcome the resistance posed on it by such a pore wall. In most soils, mechanical impedance increases with soil drying (Gao *et al*., 2016; Grzesiak *et al*., 2013; Whalley *et al*., 2005; Whitmore and Whalley, 2009). Thus alternate wetting and drying of soil can therefore temporally impede roots depending on soil matric potential.

Root adaptions to mechanical impedance encompass several strategies. Root tip phenes such as increased production of mucilage and root cap cell sloughing lubricate the root-soil interface (Boeuf-Tremblay *et al*., 1995; Iijima *et al*., 2000, 2004). Sharper root tip shape reduces stress at the root tip via a more cylindrical cavity expansion (Bengough *et al*., 2011; Colombi *et al*., 2017a). Architectural phenes, such as steeper root angles might reduce deflection upon encountering a strong layer (Dexter and Hewitt, 1978). Other phenes such as the presence of root hairs help root tip penetration by anchoring the root into the soil (Bengough *et al*., 2016). A comprehensive review of root morphological adaptions to mechanical impedance by Potocka and Szymanowska-Pułka (2018) concluded that adaptations to mechanical impedance are present across different architectural and anatomical scales. However, it is clear that limited research has been carried out discriminating root anatomical responses among root types in response to mechanical impedance.

Root anatomical variation among maize genotypes is better able to predict penetration of strong wax layers than root diameter alone (Chimungu *et al*., 2015). Mechanical impedance generally causes radial thickening of roots, including that of maize which we studied here (Bengough and Mullins, 1991; Konôpka *et al*., 2009; Materechera *et al*., 1991; Moss *et al*., 1988). This form of radial expansion is different from that resulting from secondary growth (Strock *et al*., 2018). Thicker roots buckle less (Clark *et al*., 2008; Whiteley *et al*., 1982), and modelling has found that radial expansion will reduce the stress from the root tip (Bengough *et al*., 2006; Kirby and Bengough, 2002) while simultaneously pushing particles out of the way so that the root can extend further (Vollsnes *et al*., 2010). Root thickening is associated with reduced elongation rates (Bengough and Mullins, 1991; Clark *et al*., 2001; Colombi et *al*., 2017; Iijima *et al*., 2007; Schmidt *et al*., 2013), which ultimately can result in reduced soil exploration. Roots that thicken in response to impedance do so by increasing the dimensions of the cortex (Atwell, 1990; Colombi *et al*., 2017) or both stele and cortical tissues (Atwell, 1988; Colombi *et al*., 2017; Hanbury and Atwell, 2005; Iijima *et al*., 2007; Wilson *et al*., 1977). These responses vary among plant species, root type, plant developmental stage and experimental conditions (Colombi and Walter, 2016). Cortical dimensions change by an increase in the size of cortical cells (Atwell, 1988; Hanbury and Atwell, 2005; Veen, 1982) or a combination of cortical cell size and cortical cell file number (Croser *et al*., 1999; Colombi *et al*., 2017; Iijima *et al*., 2007). Cortical cells increase their size radially, facilitated by the loosening of cell walls by microfibril reorientation (Iijima *et al*., 2007; Veen, 1982). The increase in radial cell area coincides with reduction of cell lengths (Atwell, 1988; Croser *et al*., 2000). How cell volume changes under mechanical impedance needs further clarification. Cortical cell length reduction could partly explain reduced elongation rates observed under mechanical impedance (Atwell, 1988). Further reduction of elongation rate could be caused by reduced cell production in the meristem (Croser *et al*., 2000). Recently root thickening has been directly linked to increased energy cost for root elongation with increasing soil penetration resistance for different wheat genotypes (Colombi *et al*., 2019). Root thickening has also been associated with an increase in the demand for oxygen (50% to 80%) for impeded lupin roots (Hanbury and Atwell, 2005). It is clear that root thickening has beneficial, as well as detrimental effects for the plant root system. There is a need to better understand the mechanism controlling radial thickening.

Ethylene biosynthesis and systems modified by ethylene are involved in stress responses and may regulate root responses to impedance (Atwell *et al*., 1988; Sarquis *et al*., 1991). Mechanical impedance alters maize root growth by promoting ethylene biosynthesis which inhibits elongation and promotes swelling (Sarquis *et al*., 1991). Impeded maize primary roots produced more ethylene and had an increased root diameter compared to nonimpeded roots (Moss *et al*., 1988; Sarquis *et al*., 1991). Mechanically impeded *Vicia faba* roots produced more ethylene compared to nonimpeded roots (Kays *et al*., 1974). Roots of 7-day old *Never ripe* (ethylene-insensitive) tomatoes formed elongated roots in a soft medium but were unable to penetrate a harder sand medium (Clark *et al*. 1999), and tomato roots treated with the ethylene action inhibitor 1-methylcyclopropene (1-MCP) were unable to penetrate a soft growing medium (Santisree *et al*., 2011). Based on the observed effects of ethylene on radial expansion and research indicating that thicker roots are more likely to penetrate hard soil, it has been assumed that ethylene production in response to mechanical impedance leads to radial expansion and improved soil penetration (Potocka & Szymanowska-Pułka, 2018). However in a study of *Eucalyptus* seedlings by Benigno *et al*. (2012), compacted soil reduced both ethylene production and elongation rates, suggesting that the link between ethylene production and reduced root growth is not straightforward.

Existing studies have generally focused on root length, branching and diameter responses to mechanical impedance (Konôpka *et al*., 2008). When root anatomy has been studied, different root axes have been compared while changes within a single root axis have rarely been considered. With few exceptions (Veen, 1982; Colombi *et al*., 2017), root anatomy has mainly been studied on primary roots (Hanbury and Atwell, 2005; Croser *et al*., 1999; Iijima *et al*., 2007; Colombi *et al*., 2019). However, different root classes can react differently to impedance (Vanhees *et al*., 2020). In this study we hypothesise that root radial expansion is negatively associated with the penetration rate of roots in compacted soil layers. Secondly, we assessed root class and genotypic differences in the ability of roots to penetrate hard soil and tested ethylene responsiveness variation in these groups. In this context we propose ethylene might function as a signal associated with thickening and suggest that prolonged production of ethylene in response to mechanical impedance can function as a ‘stop’ signal for axial growth of that particular root axis. Genotypes that produce less ethylene, or that are insensitive to ethylene could therefore maintain root elongation rate more easily under impeded conditions.

## Materials and methods

### Experiment 1: Anatomical changes to a root axis crossing a compacted soil layer

#### Experimental set-up

A brown earth soil (FAO classification: Stagno Gleyic Luvisol) with sandy loam texture (2% clay, 21% silt, 77% sand) was procured from local sugar beet farms through British Sugar in Newark (UK). The soil was obtained from sugar beet during the manufacturing process. Before column packing the soil was air-dried and sieved to <2 mm. Dried soil was wet to 17% gravimetric moisture content. Columns (14.8 cm diameter, 23 cm total height) were uniformly packed creating three regions with a compacted layer (1.5 g/cm^³^ and thickness of 3 cm) placed between low bulk density layers (1.2 g/cm^³^). The top and bottom areas were 7.5 and 9.5 cm long respectively, making up a total of 20 cm of total height of soil in column. A mould was used to create the compacted layer after which it was transferred onto the bottom half of the column. The soil surface of the compacted layer was abraded at each side to assure the compacted layer and the non-compacted soil above and below the compacted layer were adequately adhered. The pots were lined with a plastic sleeve to facilitate removal of the intact soil column after scanning. A preliminary trial was conducted to optimise the positioning of the compacted layer and to identify the preferred number of growing days (to account for growth up to node 4 reaching below the compacted layer).

Smaller columns (10 cm high, 5 cm diameter) packed at the same moisture content and density as the layered system were used to record penetrometer resistance and measurements were made with an Instron (Instron 5969, 50kN load cell, Instron, Norwood, USA) fitted with a penetrometer needle (0.996 mm cone diameter, 15° semi-angle). The penetrometer tip penetrated the samples for 12 mm at a constant speed of 4 mm sec^-1^. Measurements were averaged between 5 – 11 mm extension. Smaller (1.2 g/cm^³^) and greater (1.5 g/cm^³^) bulk densities had penetrometer resistance of 0.48 ± 0.03 (sd) MPa and 0.83 ± 0.01 (sd) MPa respectively and were significantly different (t-test, p = 0.002).

#### Plant material and growing conditions

Four genotypes (recombinant inbred lines; IBM086, IBM146, IBM014 and OhW128) previously studied in field trials (Vanhees *et al*., 2020; Chimungu *et al*., 2015), were selected based on their contrasting ability to penetrate the compacted layer and with sufficiently steep root angle to allow for roots to reach the compacted layer. Seeds were acquired from Dr. Shawn Kaeppler (the University of Wisconsin, Madison WI, USA – Genetics Cooperative Stock Center, Urbana, IL, USA). Seeds were sterilised (6% NaOCl in H_2_O) for 30 minutes, imbibed for 24 hours and germinated at 26 °C for 3 days before planting. Germinated seeds with similar primary root length (± 1 cm) were selected for planting. Two seeds per pot were planted 0.5 cm deep for each genotype, plants were thinned to one plant per pot if both of the seeds developed. Five blocks staggered in time were planted with one replicate for each genotype per block. Plants were grown in a greenhouse at a 25/18°C day/night temperature and a 14h/10h day/night cycle provided by additional lighting at a maximum of 600 µmoles photons m^-2^ s^-1^. Once a week a nutrient solution (100 g of HortiMix Standard: NPK ratio 15-7-30 to 1L of solution contains 107 mmole of total water soluble N, 4.5mmoles P_2_O_5_ (w/w), 32 mmoles total K_2_O (w/w), 4 mmoles MgO (w/w), 0.04 mmoles Fe-EDTA, 0.18 mmoles Mn, 0.28 mmoles B, 0.04 mmoles Zn, 0.03 mmoles Cu, 0.013 mmoles Mo (Hortifeeds, Lincoln, UK) was added when watering. Moisture content of the pots was maintained at 17% gravimetric moisture content by watering a constant amount of water per block based on the overall starting reference weight of the pots. Plants were grown for 49 days to assure sufficient growth of node 3 and node 4 roots. These nodes were selected because node 1 and 2 were too horizontally oriented to sufficiently interact with the compacted layer (more horizontal growth of earliest nodes has also been described by Araki *et al*., 2000; York *et al*., 2015).

#### X-ray Computed Tomography

Soil columns were not watered 48 hours prior to scanning to allow for enhanced contrast between the roots and soil matrix. Each column was imaged using a v│tome│x L (GE Measurement and Control Solutions, Wunstrof, Germany) X-ray µCT scanner. Two scans (multiscan option) were taken per column (top and bottom) with a total scanning time of two hours per column. The distance from the centre of the sample to the detector was 2000 mm. X-ray energy was set at 290 kV and the current was 2700 µA. Filters were fitted to the X-ray gun (1.5 mm copper, 0.5 tin) and detector (0.5 mm copper) to enhance the image quality. Image averaging was set at 5 images. The scanning resolution was 96 µm and 2400 image projections were taken for each scan.

#### Image processing and analysis

Images were reconstructed at 32-bit (Phoenix DatoS│x2 reconstruction tool, GE Sensing & Inspection Technologies GmbH, Wunstorf, Germany) with scan optimisation and beam hardening correction set at 8. The 3D image volumes were analysed in VGStudioMax 2.3 (Volume graphics Gmb, Heidelberg, Germany). The greyscale values of the two obtained volumes were equalised and scans were aligned and stitched together. An example of a scan can be found in Figure 1. Nodes 1 to 4 were identified manually from 2D projections of the scans (Figure S1). Each plant was marked at the base of the stem with a thumbnail pressed into the stem prior to scanning which served as a reference point for labelling of each root axis (Figure 1A). For each node, all roots were labelled clockwise (observed from above, yz-projection plane) around the reference point. After labelling each root axis the polyline tool within VGStudioMAX was used to trace the roots from the root base downwards (Figure 1A). Polylining stopped either at the root tip or alternatively when the column wall or bottom of the column was reached. Whether roots reached and subsequently crossed the compacted layer was recorded. Distances along the root axis were measured during polylining to determine sectioning positions relative to the compacted layer along penetrating roots. Three sectioning points were located along each selected penetrating root axis; ‘before’, 1 cm above the compacted layer, ‘within’, 1 cm after penetrating the compacted layer and ‘after’, 1 cm after crossing the layer (Figure 1B). The polylines were also used for measuring root angle and rooting depth with PAM (Polyline Analysis Measurement Software, University of Nottingham, UK), an in-house software developed for these measurements to calculate root angle from the horizontal. Separate shorter polylines were drawn right above the compacted layer, tracing the root upward over a distance of 2 cm, to determine the angle at which the roots encounter the compacted layer (Figure S2). Rooting depth per pot was taken as the average maximum depth of all roots up till their root tip or when they hit the pot wall.

**Figure 1.**
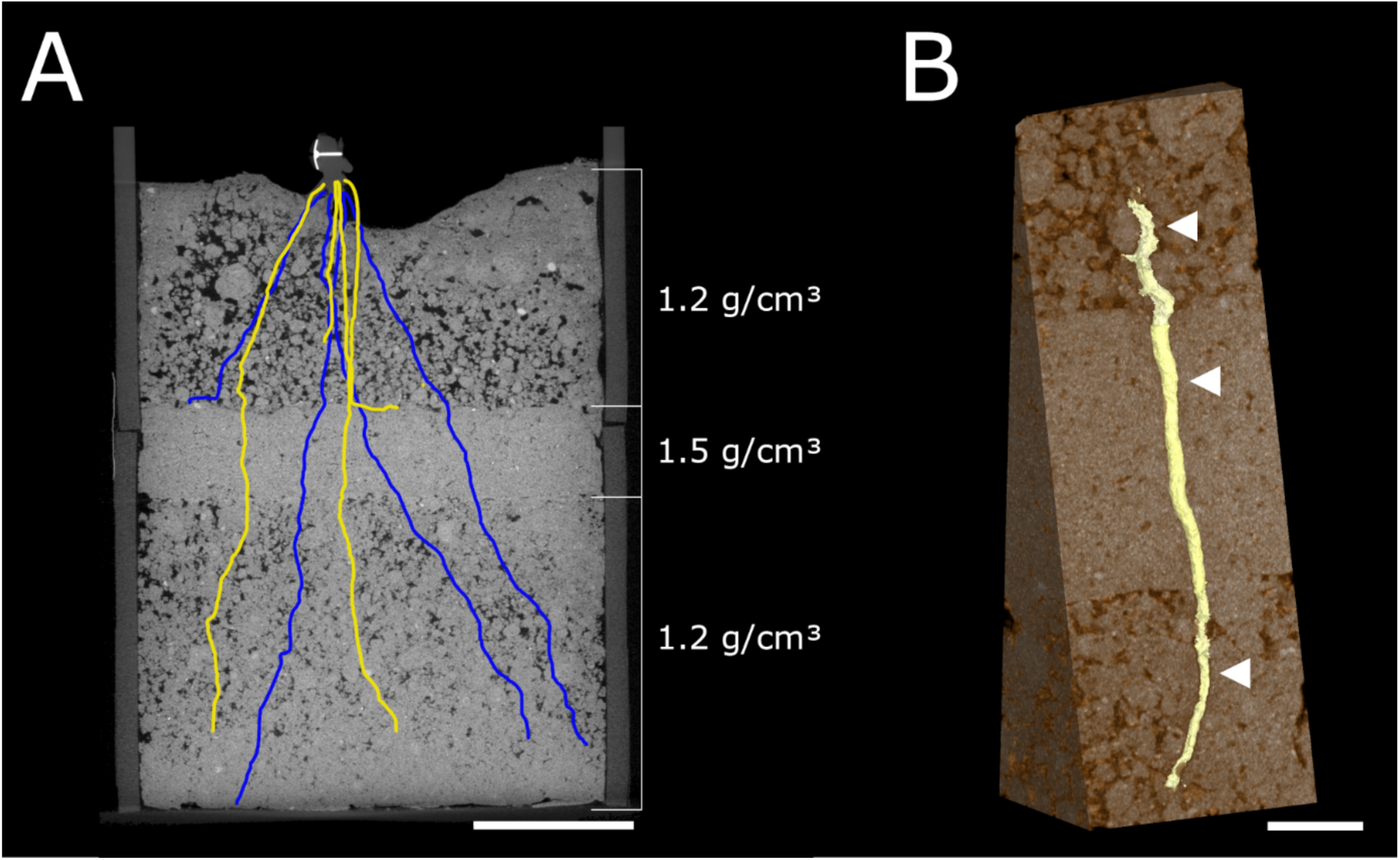
X-ray CT images/reconstruction of (A) a root system encountering a compacted layer and (B) a root growing through the compacted layer. (A) Cross sectional view of a soil column in the xy-plane with a compacted layer in between less dense layers. Blue and yellow lines represent the projection of the different polylines on the xy-plane. Colours: yellow - node 4 and blue - node 3. Scale bar at 5 cm. (B) A 3D reconstruction of a segmented root growing through the denser layer. The white arrows represent the sectioning positions along the root axis (1 cm before, within and after the compacted layer). Scale bar at 1 cm.

#### Root harvest and sectioning for root anatomical phenes

Immediately after scanning, all soil columns were lifted out of the plastic columns and roots were washed from the soil. The entire root system was extracted and stored in 75% ethanol (v/v) until sectioning. Penetrating roots of node 3 and node 4 were selected for sectioning based on polylining results and clipped from the entire root system. The length along each individual root axis was measured and sectioning positions were identified along the root axis of interest (Figure 1). Pieces of root containing the sectioning positions were excised out of the root axis and embedded by placing them into 3D printed moulds (Atkinson and Wells, 2017). 6% agarose (Sigma-Aldrich Co. Ltd, Gillingham, UK) at 39°C was used to fix the roots within the mould. A vibrating microtome (7000 smz-2) (Campden Instruments Ltd., Loughborough, UK) was used to section the roots within the agarose block at 200-230 µm thickness per slice (blade speed at 1.75-2 mm/s, blade frequency at 70 µm). Root sections were then incubated in calcofluor white (Sigma-Aldrich, Co. Lt, Gillingham, UK), 0.3 mg/ml for 90 seconds, rinsed with deionised water and placed on a microscopy slide and covered by a coverslip. Cross sectional images (Figure 2) were obtained by using an Eclipse Ti CLSM confocal scanning microscope (Nikon Instruments Europe B.V., Amsterdam, The Netherlands) with three excitation lasers. Images were collected using 10x objective, all three image channels were combined. As entire cross sections did not fit the 10x objective image space, multiple images per root section were obtained, taking care that part of each set of images overlapped. ICE software (Microsoft, Redmond, WA, US) was used to obtain one composite image per root section (camera motion set at planar motion). Image analysis for root anatomical phenes was conducted by creating object directories in objectJ (Vischer and Nastase, 2009), a Fiji plug in (Schindelin *et al*., 2012) according to Vanhees *et al*. (2020) with an additional directory for xylem vessel area. Abbreviations of root anatomical phenes can be found in Table 1.

**Figure 2.**
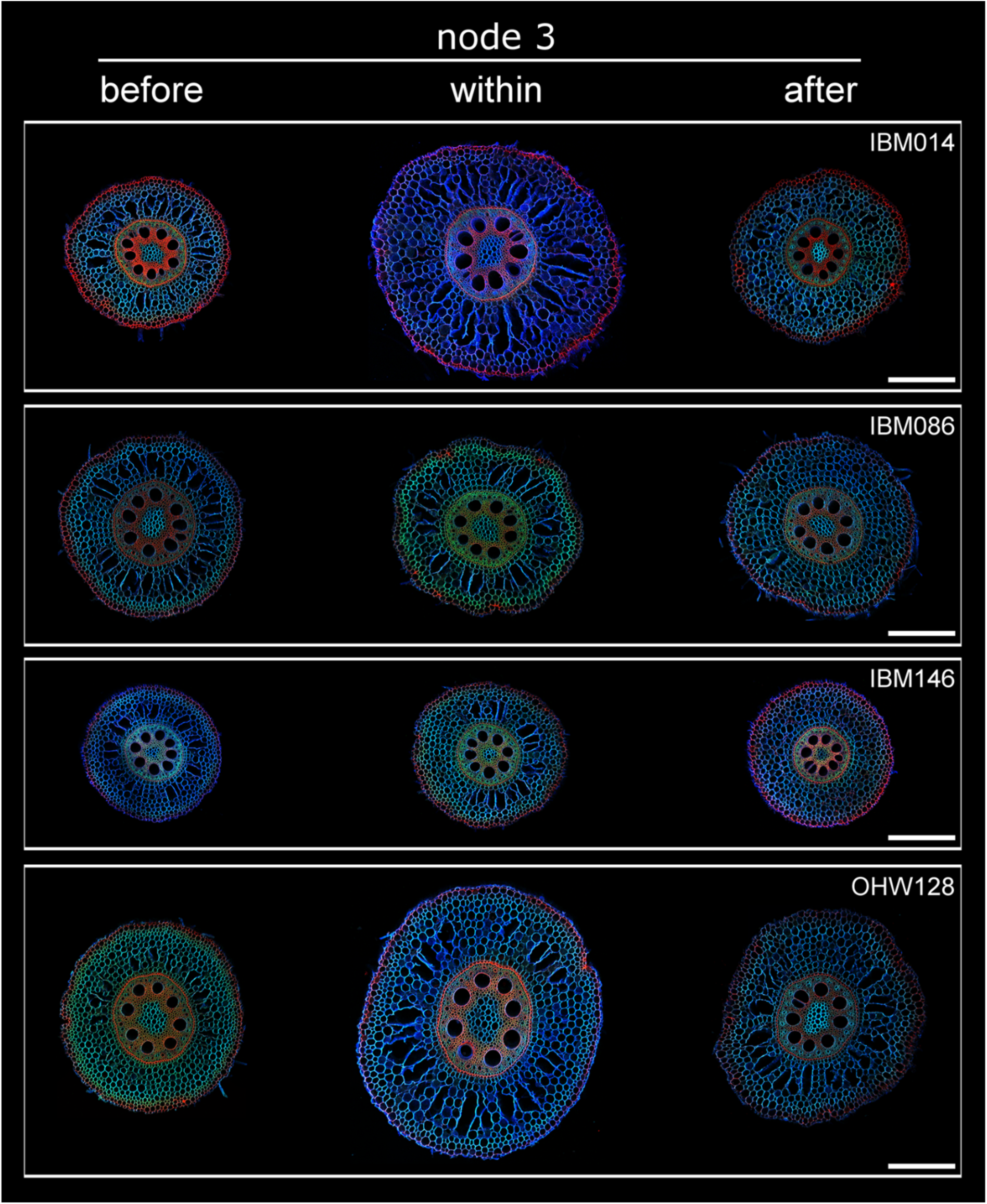
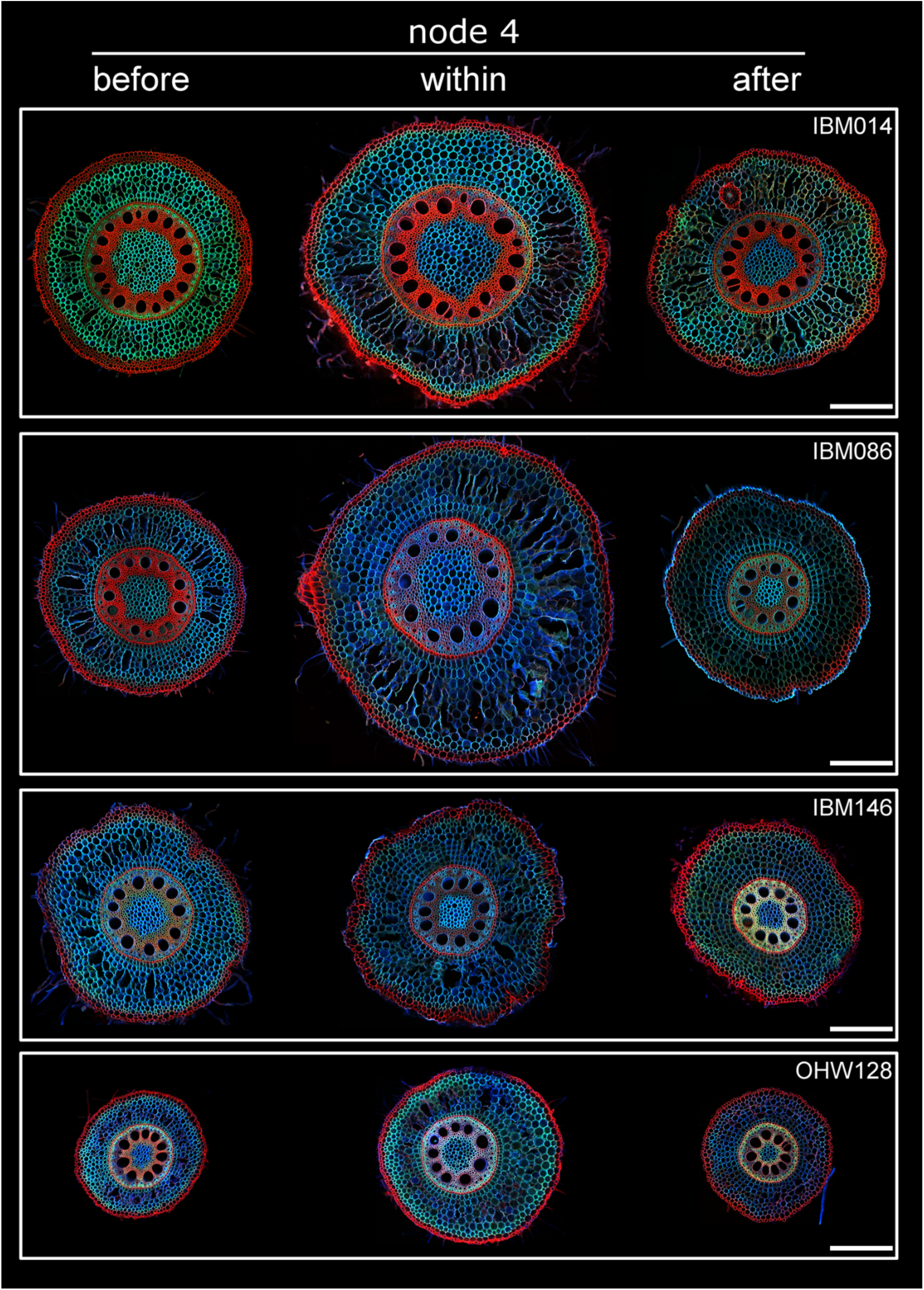
Typical images of sections taken along the same root axis from node 3 and node 4 (see continued figure) for each genotype. Before, within and after indicate the root axis position where the roots were sectioned in relation to the compacted layer. All images are at the same scale, scale bar at 500 µm.

**Table 1.** Root anatomical phenes and their abbreviations. All phenes were measured according to Vanhees *et al*. (2020).

| Abbreviation | Root anatomical phenes | Unit |
| --- | --- | --- |
| RCSA | Root cross sectional area | mm<sup>2</sup> |
| TSA | Total stele area | mm<sup>2</sup> |
| TCA | Total cortical area | mm<sup>2</sup> |
| CF | Cell file number | - |
| IN | Cell size - inner cortical region | µm<sup>2</sup> |
| MID | Cell size - middle cortical region | µm<sup>2</sup> |
| OUT | Cell size - outer cortical region | µm<sup>2</sup> |

### Experiment 2: Radial expansion is driven by ethylene

#### Plant material and growing conditions

Seeds from four genotypes (IBM086, IBM146, IBM014 and OhW128) were surface sterilized in 3% DI water in sodium hypochlorite (v/v), rolled into tubes of germination paper (76 lb, Anchor Paper, St. Paul, MN, USA), and placed in a dark chamber at 28 °C for 4 days in beakers containing 0.5 mM CaSO4. Beakers containing germinating seedlings were placed under a fluorescent light (350 µE m^-2^s^-1^) at 28 °C for one day before transplanting to an aerated solution culture. Three randomly assigned seedlings from each genotype were transplanted in foam plugs suspended above each 38 L solution culture tank. The solution culture tank contained per litre: 3 mmol KNO_3_, 2 mmol Ca(NO_3_)_2_, 1 mmol (NH_4_)_2_HPO_4_, 0.5 mmol MgSO_4_, 50 mmol Fe-EDTA, 50 mmol KCl, 25 mmol H_3_BO_3_, 2 mmol MnSO_4_, 2 mmol ZnSO_4_, 0.5 mmol CuSO_4_ and 0.5 mmol (NH_4_)_6_Mo_7_O_24_. The pH was adjusted daily to 5.5 using KOH and the solution was completely replaced every 7 days. Plants were grown for 30 days in a climate chamber. During the growth period, the mean minimum and maximum air temperatures were 26 ± 3°C and 30 ± 3°C, respectively with maximum illumination of 800 μmol photons m^-2^ s^-1^ and average relative humidity of 40%.

#### Ethylene application

Three replicates of all four genotypes (i.e. each 38 L tank) were exposed to one of four different treatments (1) root zone air application (control), (2) root zone ethylene application (dose 1), (3) root zone ethylene application (dose 2) and (4) root zone 1-MCP (1-methylcyclopropene, ethylene inhibitor) application, all applied continuously beginning at seedling transfer to solution culture. Solution culture tanks in the control treatment were bubbled at 10 mL min^-1^ with ambient air in 38 L of solution culture. In the ethylene treatment (dose 1), compressed ethylene (1 mL L^-1^ in air, as used by (Gunawardena *et al*., 2001)) was bubbled through 38 L of solution culture at 10 mL min^-1^. In the ethylene treatment (dose 2), compressed ethylene (1 mL L^-1^ in air) was bubbled through 38 L of solution culture at 20 mL min^-1^. For the 1-MCP treatment, 1-MCP (SmartFresh, ∼3.8 % active ingredient, AgroFresh, USA) was volatilized by dissolving 0.17 g in 5 mL water in a glass scintillation vial, and then transferred into a 2-L sidearm flask. An open-cell foam plug enclosed the mouth of the flask, and the headspace containing 1-MCP gas was bubbled through 38 L of solution culture at a rate of 10 mL min-1. The air pump ran continuously, and the 1-MCP was replenished daily into the sidearm flask. There was no significant effect of flow rate on headspace ethylene concentrations, which ranged from 0.78-1.58 μL L^-1^ with a mean of 1.15 μL L^-1^, therefore the results of ethylene treatments were combined in a single mean. After 30 days of growth, plants were sampled. Third and fourth whorl nodal roots from each plant were sampled 5-8 cm from the base of the plant and preserved in 75% EtOH (v/v) for further anatomical analysis.

#### Laser Ablation Tomography and evaluation of root anatomy

Root anatomy was imaged using Laser Ablation Tomography (LAT) (Hall *et al*., 2019; Strock *et al*., 2019) In brief, a pulsed UV laser is used to vaporize the sample at the camera focal plane and simultaneously imaged. Imaging of root cross-sections was performed using a Canon T3i camera (Canon Inc. Tokyo, Japan) and 5× micro lens (MP-E 65 mm). Two images for each root sampled were collected for phenotypic analysis. Six anatomical phenes (Table 1) on every image were measured using objectJ (Vischer and Nastase, 2009) and a Fiji plug in (Schindelin *et al*., 2012) according to Vanhees *et al*. (2020).

### Statistical analysis

For experiment 1 the number of replicates obtained per genotype and node varied as one plant (genotype OhW128) died during the 49 day growth period. Hence for node 3 and 4 only four replicates were taken into account for this genotype. For genotype IBM014, node 4 roots were underdeveloped (<0.5 cm long, observed during washing) at sampling, therefore we only obtained four replicates for this measurement. Additionally, not all genotypes were equal in crossing the compacted layer, hence some genotypes have fewer replicates at the within and after the compacted layer sectioning positions. Both the effect of blocking and interaction effects were tested, when not significant they were omitted from the analysis. Factorial regression was used to assess the effect of different factors on root counts. A Poisson distribution was used followed with *post-hoc* Tukey comparisons to compare factor levels. Correlations between root angle and count data were calculated using a Spearman-Rank correlation. Penetration rates were calculated per node as the ratio of roots that crossed the layer and reached the layer. An ANCOVA was performed to assess the relationship between root angle 2 cm above the compacted layer, genotype and penetration rate. Root thickening was defined as the increase of overall root cross sectional area and an ANOVA was used to identify the effect of factors genotype and node. Anatomical changes were similarly assessed by ANOVA that included factors genotype, node and sectioning position on root cross sectional area, total stele area, total cortical area and cell file number. The same factors were used with the addition of the cortical region for the ANOVA on cell size. Tukey comparisons were carried out between nodes, between genotypes within nodes and between sectioning positions for root cross sectional area. For cortical cell size and cell file number Tukey comparisons were used to identify differences between sectioning positions. The increase of cell size was calculated for the different cortical regions and for the different nodes. For experiment 2 average cortical area, stele area and cell file number were assessed by ANOVA and Tukey comparison identified differences between ethylene, 1-MCP and control treatments. Root anatomical measurements were compared between the two experiments and differences across treatments were assessed by Tukey comparison. Correlations between cortical cell size obtained from both experiments were calculated.

## Results

### Experiment 1: Anatomical changes within a root axis crossing a compacted layer

#### Steeper roots were more likely to reach the compacted layer

Although the same number of roots was formed per node irrespective of genotype or node (Figure 3A, Table 2) the number of roots reaching the compacted layer varied among genotypes. Within a node, the number of roots reaching the compacted layer was not different among genotypes (Figure 3A). However, significantly fewer roots reached the layer for node 3 roots of genotype IBM086 in comparison with node 4 roots of genotype IBM146 (Figure 3A). The number of roots reaching the layer was only significantly different from the number of roots crossing the layer for node 4 roots of IBM086 (Figure 3B). Younger nodes (node 4) were steeper than older nodes (node 3) (Figure 4A) and root angle was correlated with the number of roots that reach the compacted layer (Spearman’s rank correlation r=0.53) (Figure 4B). Root angle itself was node and genotype dependent (Table 2B) and steeper root angle was associated with improved penetration rates (Figure 4C). IBM086 had the most shallow-angled roots (Figure 4A), which led to node 3 roots hitting the pot-wall before reaching the compacted layer.

**Figure 3.**
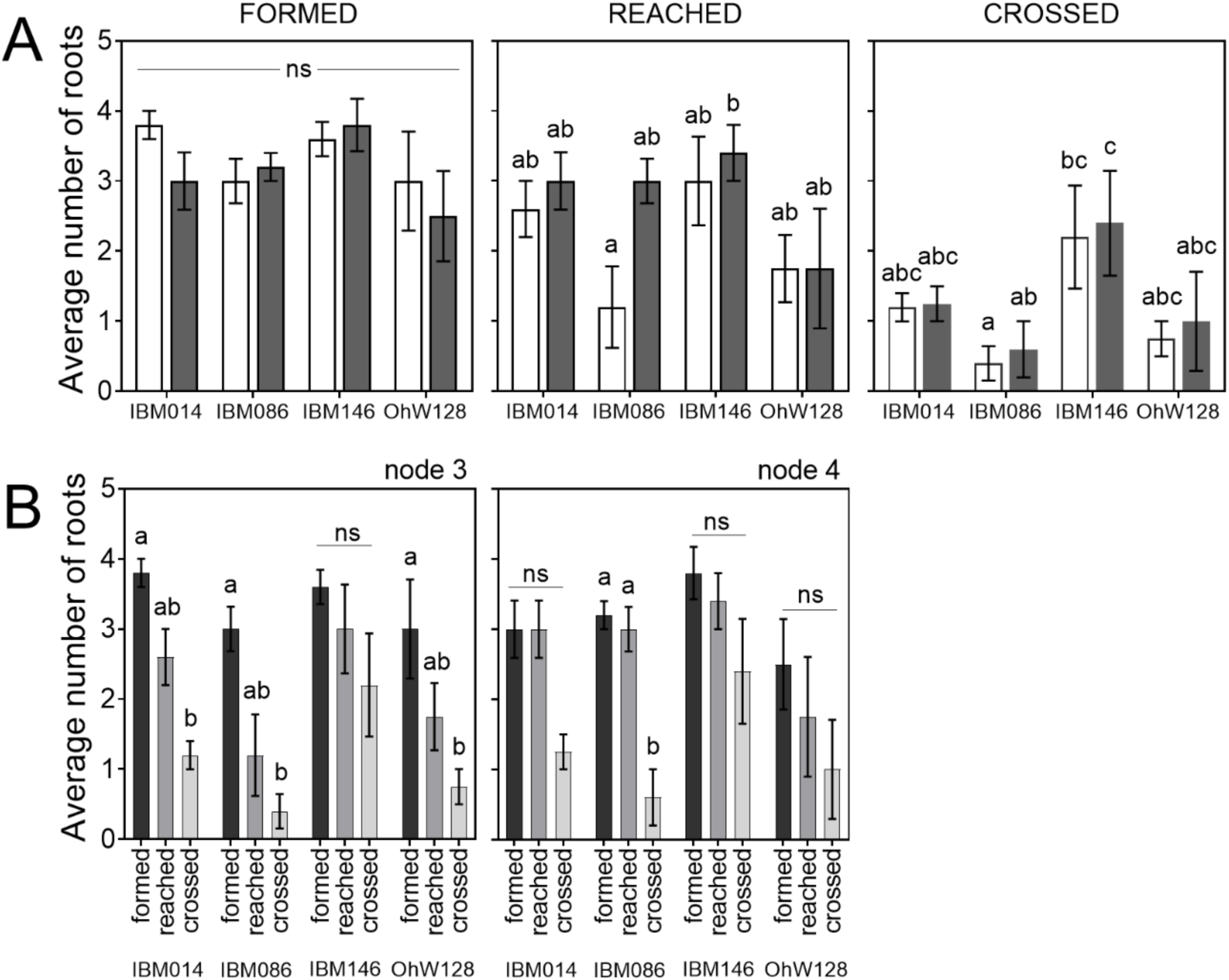
(A) Root counts at different locations with respect to the compacted layer. Bars in white are root counts for node 3, bars in grey are root counts for node 4. Differences in root counts between nodes and genotypes were assessed with Tukey comparisons (P ≤ 0.05). (B) Root counts per node and genotype on different locations with respect to the compacted layer. Differences between root counts are shown by different letters, based on a Tukey comparison (p ≤ 0.05) within node and genotype combinations. ns stands for non-significant.

**Figure 4.**
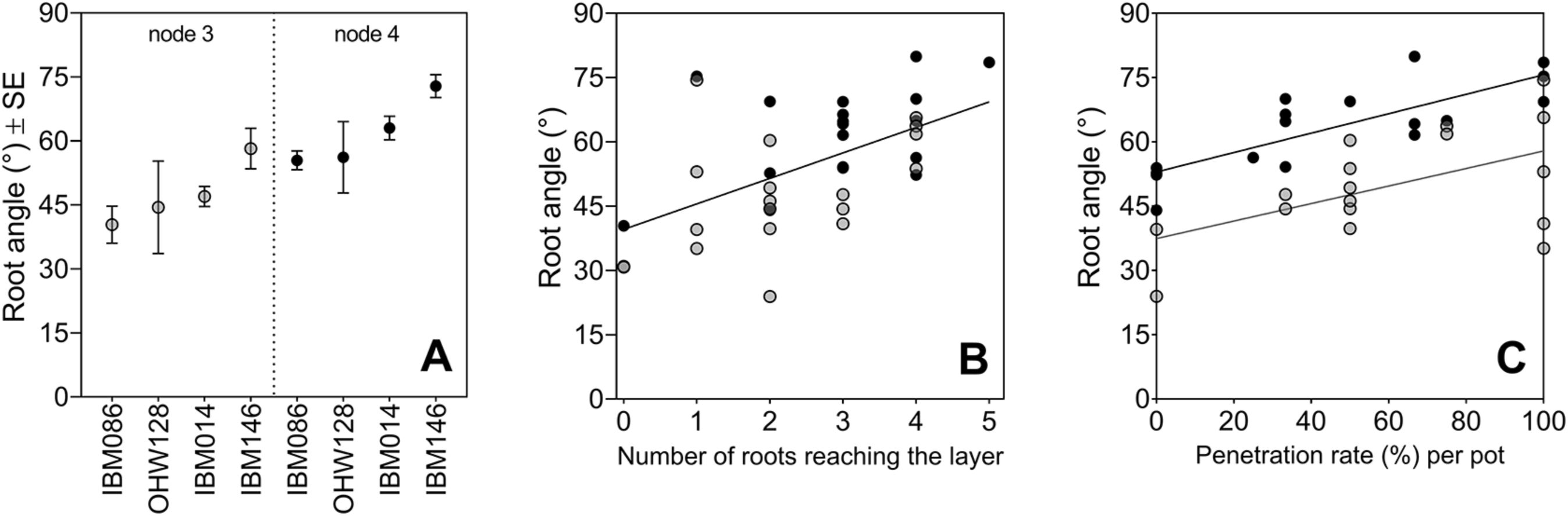
Root angle is different between nodes and determines if roots reach the compacted layer, with steeper roots having greater penetration rates (A) Mean ± SE for different genotypes per node. (B) Correlation between root angle and the number of roots reaching the layer. Correlations were tested with a Spearman rank correlation (r=0.5318, p=0.0007). (C) Linear relationships between root angle measured at the crown and the penetration rate for each pot in the study. Significant R² values of 0.28 (p=0.0269) and 0.64 (p<0.0001) for node 3 and node 4 respectively. For all figure panels node 3 data is visualised in grey and node 4 data in black.

**Table 2.** (A) Factorial regression for total number of roots and (B) root angle for node 3 and 4 roots. The variable ‘position’ refers to the number of roots counted before the compacted layer, within the compacted layer and after the compacted layer. Significance at ** p ≤ 0.01 and *** p ≤ 0.001.

| Total number of roots |  |  |  |  |
| --- | --- | --- | --- | --- |
| A | Factor | Deviance | p (> Chi) |  |
|  | Position | 35.47 | 1.99E-08 | *** |
|  | Genotype | 12.40 | 6.14E-03 | ** |
|  | Node | 0.80 | 0.44 |  |
| Root angle |  |  |  |  |
| B | Factor | F-value | p-value |  |
|  | Genotype | 5.39 | 4.06E-03 | *** |
|  | Node | 17.45 | 2.12E-04 | ** |

#### Genotypes differed in their ability to penetrate a compacted soil layer

The number of roots crossing the compacted layer varied among genotypes (Figure 3A). IBM146 had more roots crossing the compacted layer (Figure 3A) in comparison with IBM086 where roots did not fully reach the compacted layer (node 3) or did not cross the compacted layer (node 4). Higher percentages of roots grew into the layer than across it (Table 3). When roots did not grow into the compacted layer, they either buckled or deflected at the layer (Figure S3). When roots buckled, swelling of the root tip was observed. Penetration percentages varied among genotypes (Table 3), and penetration rate was greater when roots were steeper at the crown (Figure 4C). No differences were found between nodes for root angle right above the layer, however steeper root angles at this position were associated with greater penetration rate (Figure 5). The average rooting depth of nodal roots depended on the node, and overall roots of node 3 were shallower than roots of node 4 (Figure 6). Roots of genotype IBM146 grew to the greatest depth for both nodes (Figure 6) and were the steepest (Figure 4A).

**Figure 5.**
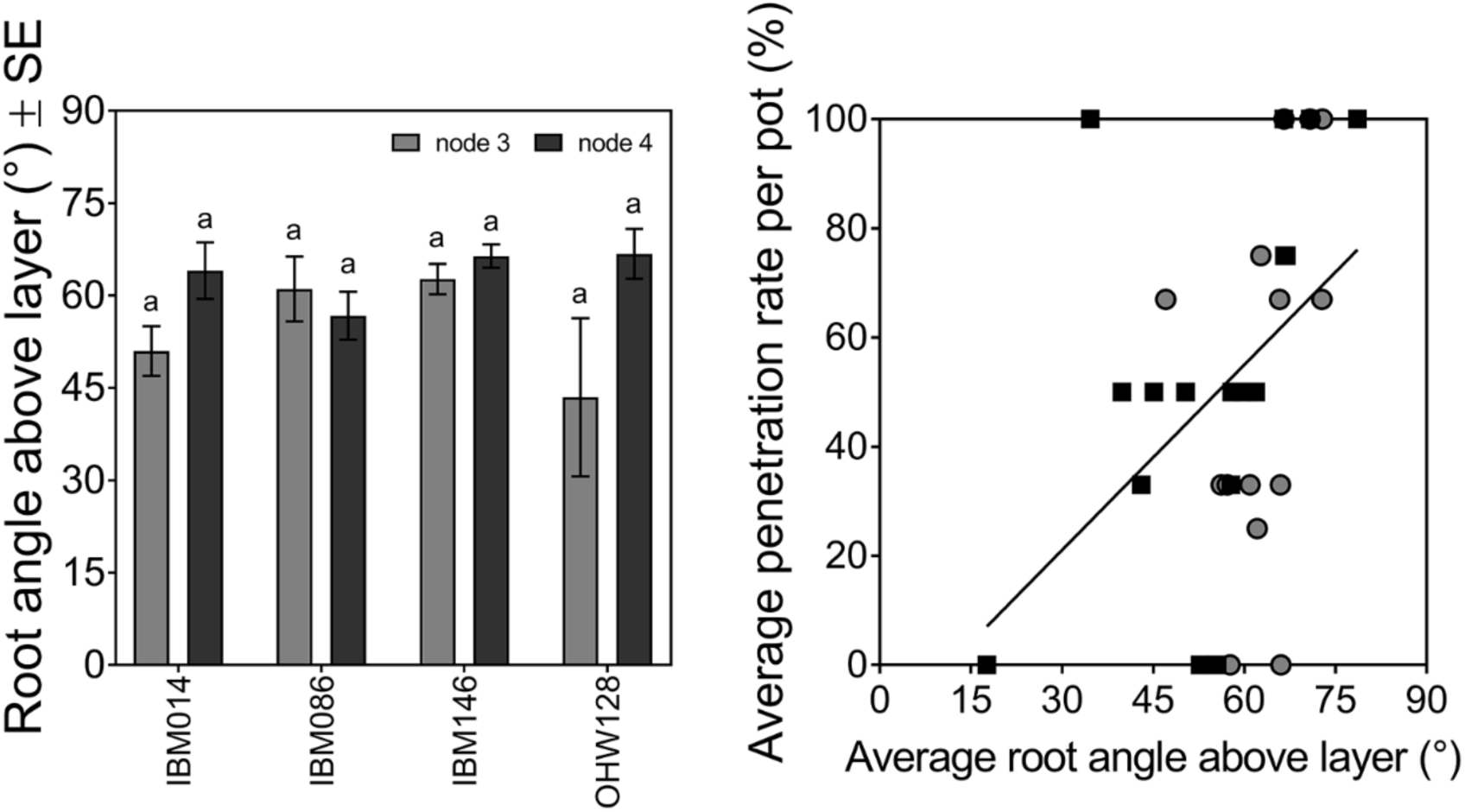
The angle at which the roots approach the layer for node 3 and node 4 is the same (Tukey comparison (p ≤ 0.05)), while root angle does influence the penetration rate per pot significantly (p=0.02, R^2^=0.25). Node 3 data in grey and node 4 data in black.

**Figure 6.**
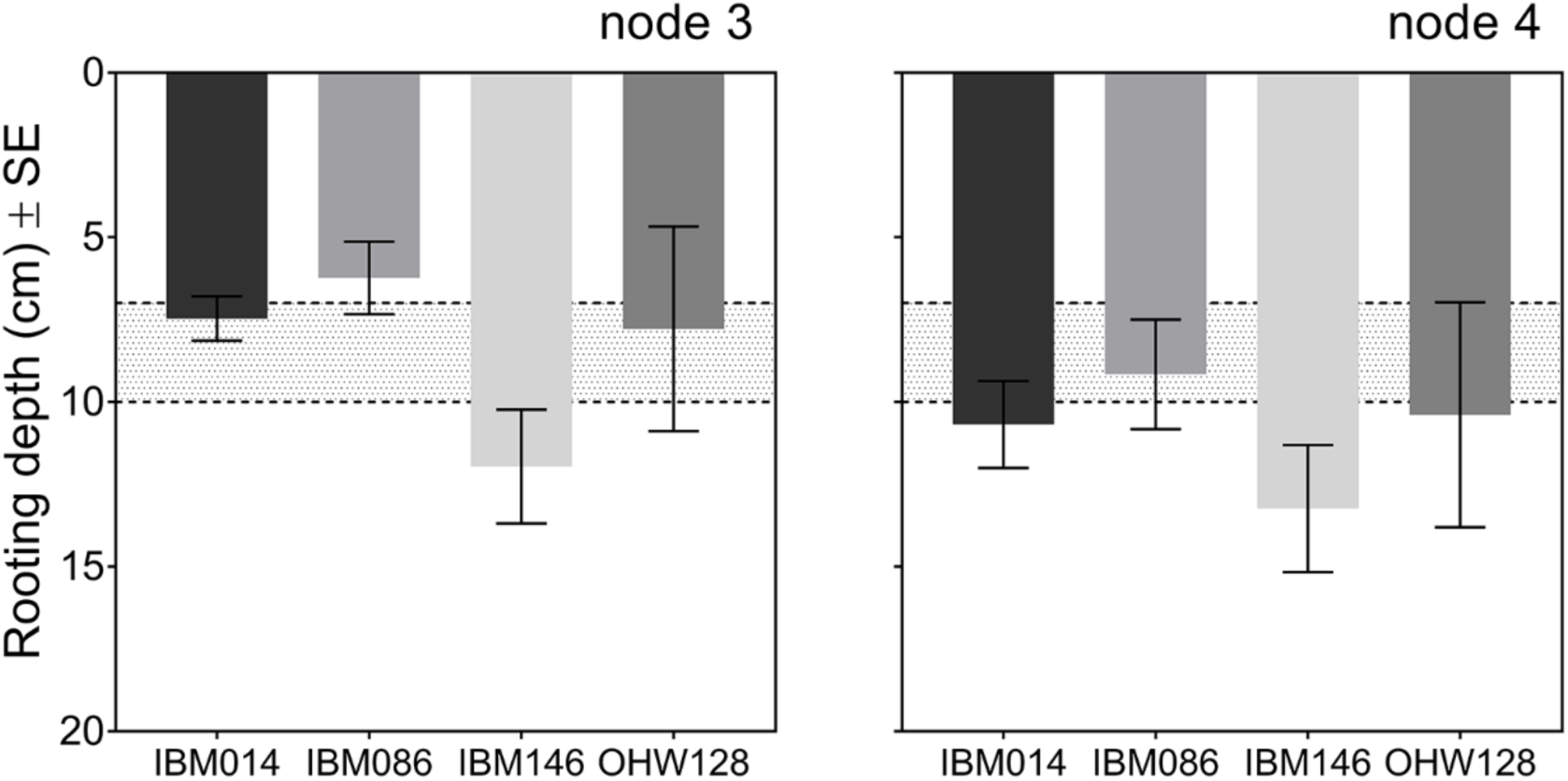
Average rooting depth (cm) ± SE per node and genotype, averaged for each replicate. Depth was calculated including all roots. If roots hit the column wall depth was recorded as the depth at which they hit the column wall. The greater bulk density layer was located at 7 – 10 cm depth and depicted by the dotted lines and grey area on the graph.

**Table 3.** Penetration rates ± SE per genotype for roots that reached the layer. Penetration rates can be seen as initially growing into the layer or roots that were able to fully cross the layer.

| Genotype | into the layer |  |  |  | across the layer |  |  |  |
| --- | --- | --- | --- | --- | --- | --- | --- | --- |
|  | Node 3 |  | Node 4 |  | Node 3 |  | Node 4 |  |
| IBM014 | 78% | ± 10 | 50% | ± 7 | 47% | ± 3 | 44% | ± 9 |
| IBM086 | 72% | ± 11 | 47% | ± 20 | 50% | ± 22 | 20% | ± 13 |
| IBM146 | 95% | ± 5 | 93% | ± 7 | 60% | ± 17 | 67% | ± 16 |
| OhW128 | 79% | ± 13 | 67% | ± 29 | 58% | ± 25 | 58% | ± 26 |

#### Radial expansion in response to impedance was dependent on genotype and nodal position

Root cross sectional area was affected by root node, genotype and sectioning position (Table 4, Figures 2, S4). The older node (node 3) had significantly smaller root cross sectional areas then the younger node (node 4) at sectioning positions before and within the compacted layer (Figure S4). However, root cross sectional areas of roots from the two nodes after crossing the compacted layers were not significantly different (Figure S4). Most genotypes thickened when crossing the compacted layer (Figures 2, 7, S4). Radial expansion was affected by genotype, node, and their interaction (Table 5). The average number of roots that crossed the compacted layer for both nodes of IBM086 and OhW128 was less than 1, hence caution should be taken interpreting thickening of these root axes. Roots from node 4 of genotype IBM014 and IBM086 thickened more than those of IBM146 (Figure S4). Thickening was absent for IBM146 node 4, since root cross sectional area from the ‘before’ and ‘within’ the compacted layer sectioning positions were not significantly different (Figure 2, S4). After roots crossed the compacted layer, root cross sectional areas returned to similar dimensions seen at the ‘before the compacted layer’ sectioning position (Figure S4).

**Table 4.** ANOVA results for root cross sectional area (RCSA), total cortical area (TCA), total stele area (TSA), cell file number (CF) and cortical cell size. Significance levels at *** p≤0.001, ** p≤0.01, * p≤0.05. ns stands for non-significant. na stands for not applicable as RCSA, TCA, TSA and CF are not associated with a specific cortical region. For cortical cell size, only the significant effects were listed. F-values and p-values can be found in Table S2.

| Factor | RCSA | TCA | TSA | CF | Cortical cell size |
| --- | --- | --- | --- | --- | --- |
| Node | *** | *** | *** | *** | ** |
| Genotype | *** | *** | *** | * | *** |
| Sectioning position | *** | *** | *** | ns | *** |
| Cortical region | na | na | na | na | *** |
| Node:Sectioning position | *** | * | ns | ns | ** |
| Genotype:Sectioning position | ns | ns | ns | ns | *** |
| Node:Genotype | ns | ns | ns | ns | ** |
| Node:Genotype:Sectioning position | ns | ns | ns | ns | * |
| Sectioning position:Cortical region | na | na | na | na | * |

**Table 5.**
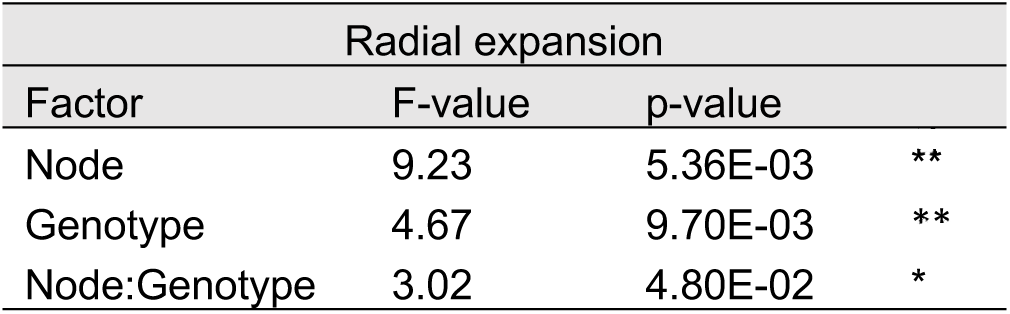
ANOVA results for radial expansion (i.e. absolute increase in root cross sectional area), measured as an increase in root cross sectional area, in response to mechanical impedance. Significance levels at *** p≤0.001, ** p≤0.01, * p≤0.05.

**Figure 7.**
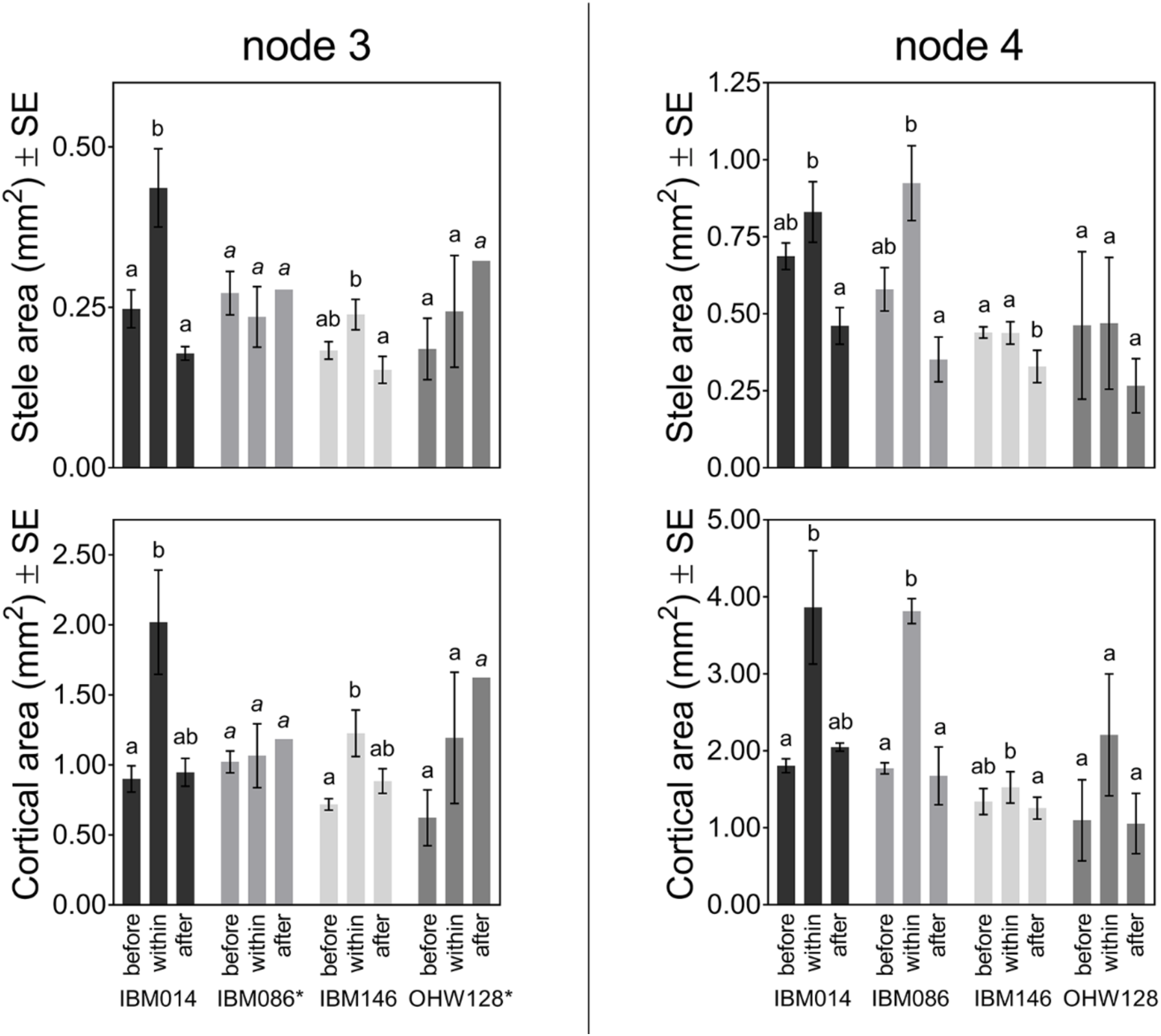
Average stele area and cortical area (± SE) at different sectioning positions (before, within and after a compacted layer) along a root axis for node 3 and 4. Differences among sectioning positions were calculated by Tukey comparisons within node - genotype combinations (P ≤ 0.05).Genotypes with * had few roots capable of crossing the compacted layer leading to a reduced number of roots that could be sectioned. Cursive letters mean separation letters indicate that replicate numbers were less for IBM086 from n=3 (before), n=2 (within) to n=1 (after) and for OhW128 from n=4 (within) to n=1 (after). When n=1 there are no SE.

#### Root thickening is more related to expansion of the cortex than the stele

Root cross sectional area, total cortical area and total stele area were dependent on node, genotype and sectioning position (Table 4). Thickening was due to increased cortical and stele areas (Figure 7, Table S1), which were correlated (Figure S5) However, there was no significant increase in stele area of node 4 roots of IBM014; this genotype thickened upon encountering the compacted layer due to cortical area increase (Figure 7). Overall the cortical tissues expanded more than the stele (Figure 7, Table S1) and the cortex has more area overall.

#### Cortical expansion is due to cellular size changes and not cell file changes

Cell size varied across the cortex (Table 4). The middle cortical cells had the largest cell sizes, surrounded by outer and inner cells with smaller cell sizes (Figure 8). Cortical cell size was also dependent on nodal position, genotype and sectioning position in relation to the compacted layer (Table 4, Figure 8). Cortical cell sizes from all cortical regions increased for those genotypes that thickened within the compacted layer (Figure 8, Table 6), while for IBM146 (node 4), there was no thickening and cell size remained constant (Figure 8). For OHW128, there was no significant increase in cell size in any part of the cortex (Figure 8). Cell sizes below the compacted layer were similar to those above the layer (Figure 8). For thickening genotypes, the outer cortical cells had a greater relative cortical cell size increase than the inner and middle cortical cells (Table 6). Despite this greater relative increase in cell size, the outer cortical cells remained smaller than the middle cortical cells at all sectioning positions (Figure 9).

**Figure 8.**
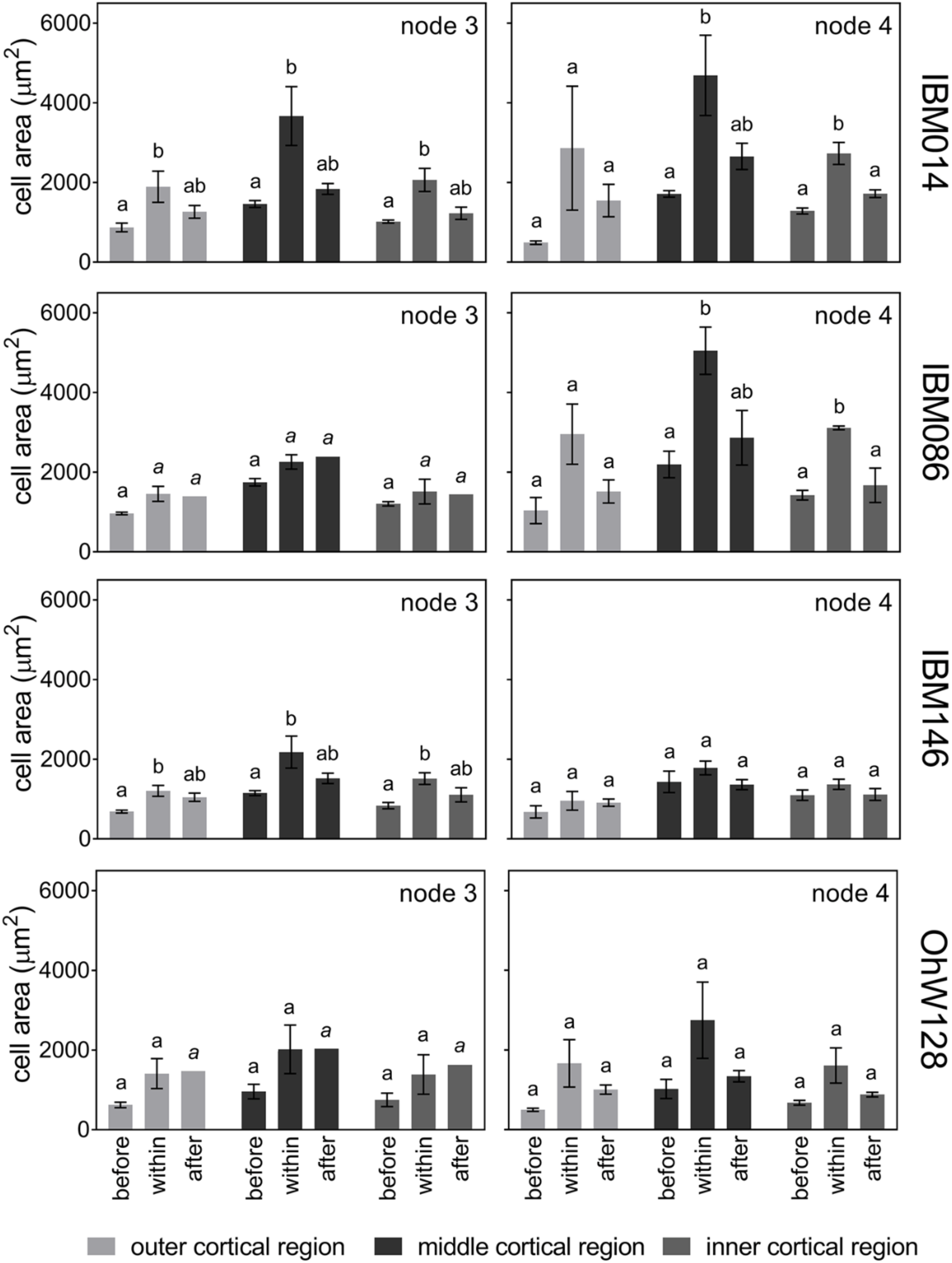
Cortical cell size (µm²) ± SE for different cortical cell regions within root cross sections. Cell size was measured along node 3 and node 4 root axes before, within and after passing the compacted layer. Differences among sectioning positions were calculated by Tukey comparisons within node - genotype combinations (P ≤ 0.05). Cursive mean separation letters indicate that replicate numbers were less for IBM086 from n=3 (before) to n=2 (within) to n=1 (after) and for OhW128 from n=4 (within) to n=1 (after). There is no SE when n=1.

**Figure 9.**
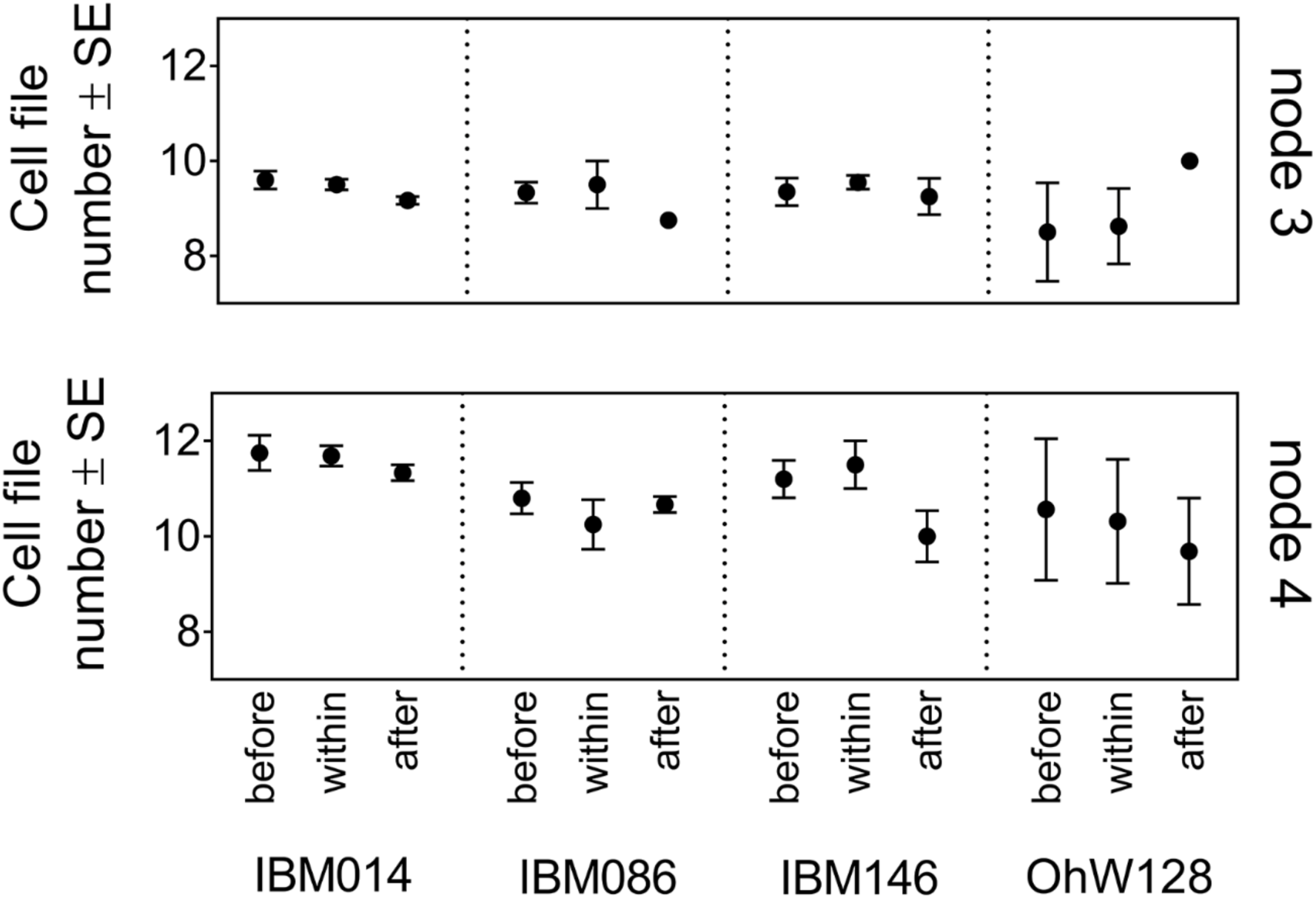
Average cell file number ± SE for different nodes and genotypes along the root axis. Cell file numbers differ between nodes. No significant differences were found among sectioning positions (before, within and after a compacted layer). There is no SE when n=1 (node3; IBM086 and OhW128).

**Table 6.** Fold increase of cell size either due to growing into the compacted layer (experiment 1) or exposure to ethylene (experiment 2). Data is depicted according to cortical area (outer, middle, inner) and genotype for node 3 and node 4.

| Experiment 1 – soil compacted layer |  |  |  |  |  |  |
| --- | --- | --- | --- | --- | --- | --- |
| Node 3 |  |  |  | Node 4 |  |  |
| genotype | outer | middle | inner | outer | middle | inner |
| IBM014 | 2.28 | 1.97 | 1.77 | 5.48 | 2.78 | 2.14 |
| IBM086 | 1.56 | 1.32 | 1.23 | 3.19 | 2.35 | 2.24 |
| IBM146 | 1.80 | 1.90 | 1.81 | 1.46 | 1.43 | 1.30 |
| OhW128 | 2.24 | 2.17 | 1.74 | 3.73 | 3.23 | 2.54 |
| Experiment 2 - hydroponics |  |  |  |  |  |  |
| Node 3 |  |  |  | Node 4 |  |  |
| genotype | outer | middle | inner | outer | middle | inner |
| IBM014 | 2.32 | 2.45 | 2.46 | 1.89 | 1.89 | 1.91 |
| IBM086 | 1.09 | 1.05 | 1.08 | 2.03 | 2.05 | 2.09 |
| IBM146 | 1.63 | 1.62 | 1.60 | 2.29 | 2.27 | 2.37 |
| OhW128 | 1.99 | 2.06 | 2.04 | 1.38 | 1.42 | 1.43 |

Cell file number was significantly different among nodes and genotypes (Table 4). Each genotype had fewer cell files for node 3 than for node 4 (Figure 9). Cell file numbers were not significantly different among sectioning positions along the root axis with respect to the compacted layer (Table 4). For all genotypes the cell file number remained stable when crossing the compacted layer (Figure 9). Therefore, radial expansion was due to increased cell size rather than increased cell file number.

### Experiment 2: Ethylene caused radial expansion

A second experiment was set up to assess the role of ethylene in radial thickening of different genotypes, different nodes and different tissues. The application of ethylene increased the cortical area in some cases but did not affect stele area (Figure 10). Genotypes varied in ethylene responsiveness, for example node 3 roots of IBM014 had the greatest increase in cortical area in comparison with node 3 roots of other genotypes (Figure 10). Roots of nodes 3 and 4 differed in their response to ethylene application, for instance in cortical area of node 3 but not node 4 roots responded significantly to ethylene application for genotypes IBM014 and IBM146 while the opposite was true for IBM086 (Figure 10). Control roots and roots treated with 1-MCP were indistinguishable for cortical and stele area (Figure 10). Since 1-MCP blocks the effect of ethylene it can be assumed that control roots were not responding to endogenous ethylene. The lack of effect was not due to inadequate concentrations of 1-MCP, since 1-MCP treated plants showed reduced root length and greater lateral branching densities in comparison with control and ethylene treatments (Figure S6).

**Figure 10.**
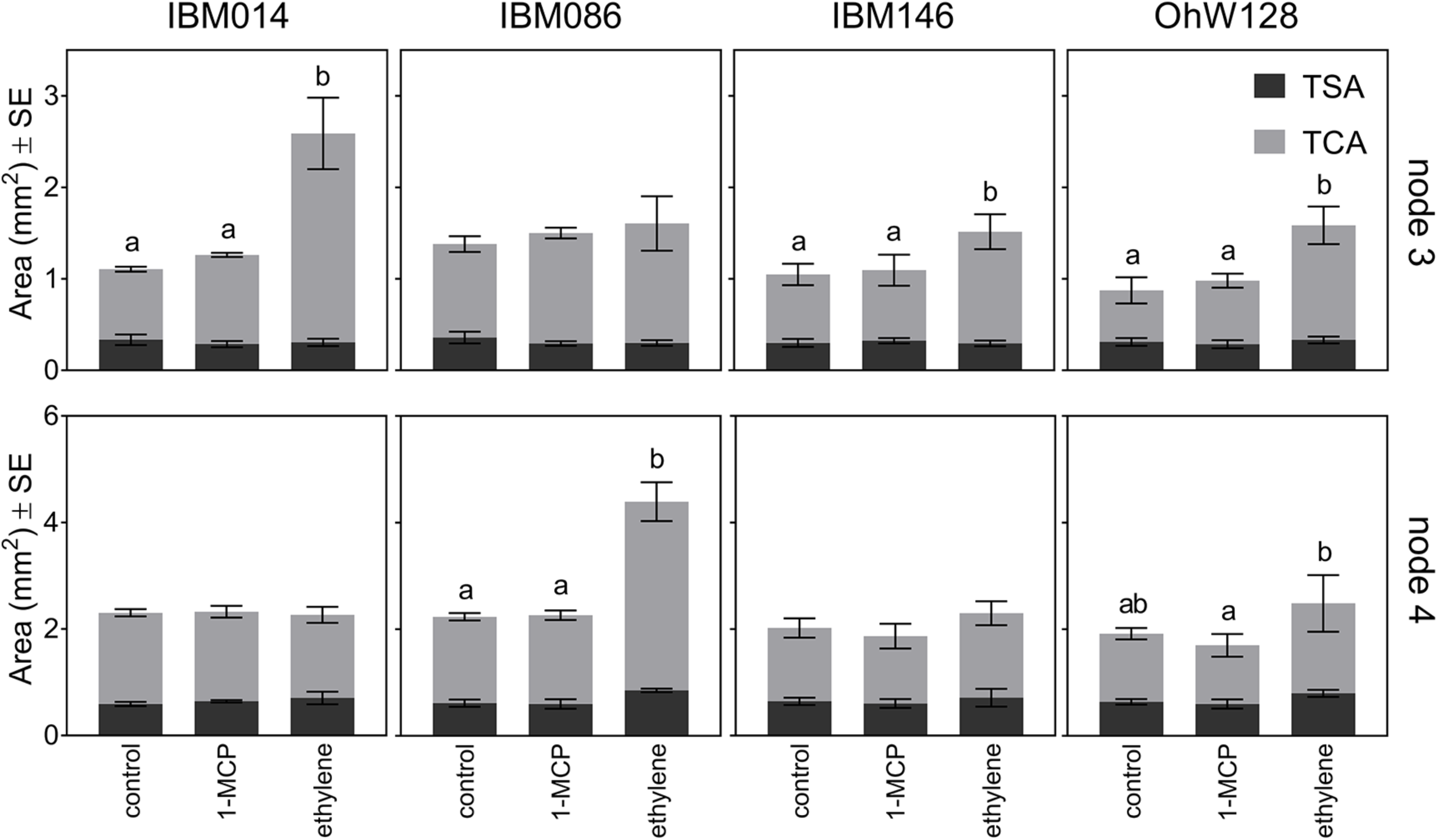
Average cortical area and stele area ± SE of root cross sections under ethylene, 1-MCP and air treatments per node and genotype. Cortical areas are shown in light grey and stele area are shown in dark grey. No significant differences were found in stele area. Lower case letters were used to identify differences among cortex areas within node and genotype according to Tukey’s test (P ≤ 0.05). Where no letters are shown, differences between treatments were non-significant;

### Comparing soil and ethylene results

Root swelling responses in independent impedance (experiment 1) and ethylene treatment (experiment 2) experiments were similar (Figures 11, 12). Root cross sectional area observed at the sectioning position before the compacted layer (experiment 1) was similar to root cross sectional area observed under control conditions in the ethylene experiment (experiment 2), across all genotypes and node combinations (Figure 11). Root cross sectional areas under impeded conditions (within the compacted layer in experiment 1) and with ethylene exposure (experiment 2) were the same with the exception of node 4 roots of IBM014 (Figure 11). The smaller root cross sectional area under ethylene can be partially due to a cell file difference of approximately 2 cell files for this genotype (Figure S7).

**Figure 11.**
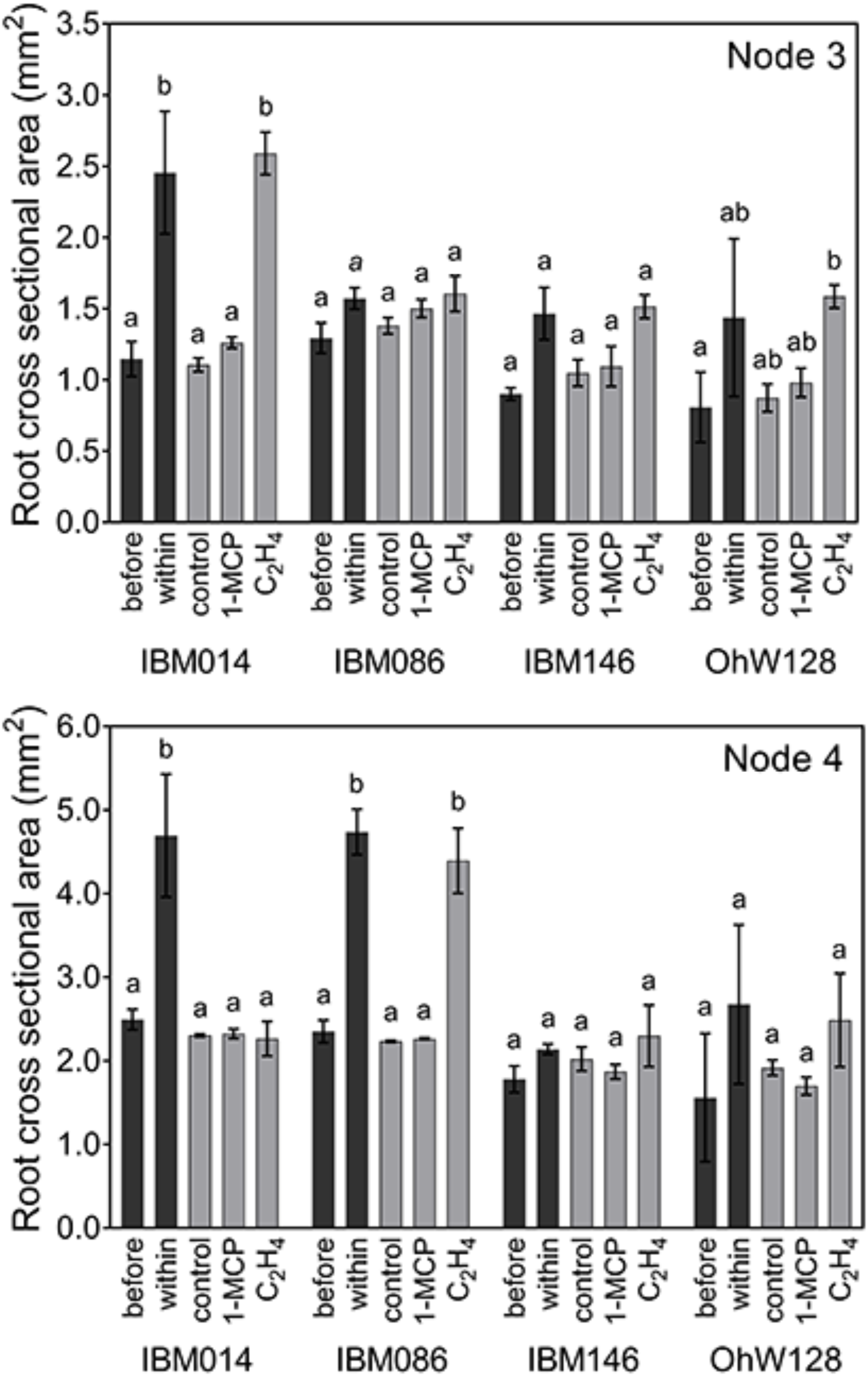
Comparison of root cross sectional area ± SE of experiment 1 (before and within compacted layer: black columns) and experiment 2 (control vs. ethylene vs. 1-MCP, grey columns) for the different genotypes and nodes. Letters show the differences between treatments assessed by Tukey comparisons within node-genotype combinations (P ≤ 0.05). Cursive mean separation letters indicate when replicate numbers dropped for IBM086 to n=2.

**Figure 12.**
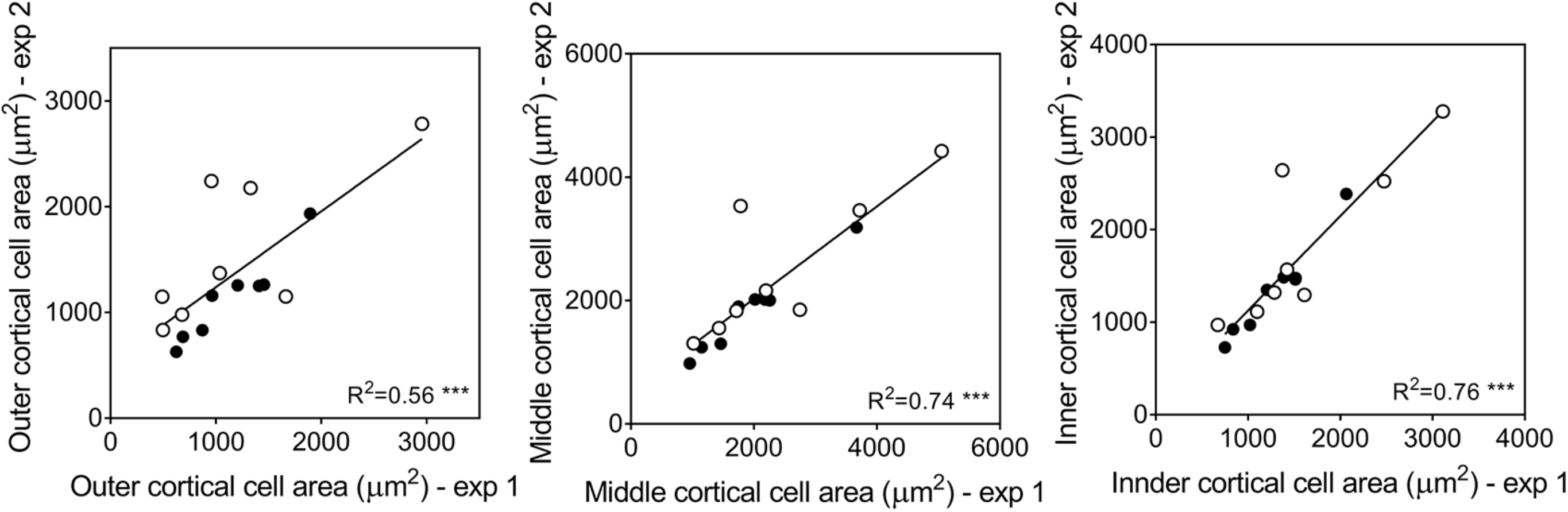
Correlation between cell size from different cortical regions of experiment 1 (pot trial in soil) and experiment 2 (grown hydroponically). Each point represents the average cell area of a genotype for paired data of ‘before the layer’ and control or paired data of ‘within the layer’ and ethylene treatment. Black circles were used for data of node 3 and white circles for data of node 4. *** level of significance at p ≤ 0.001.

When ethylene was applied, most roots thickened (Figure 11), with the following three exceptions: 1) Genotype OhW128 had greater variance, which made the increase in root cross sectional area non-significant for node 3 in soil, while for node 4 ethylene application did not cause thickening; 2) For IBM086 no thickening was observed in response to ethylene for node 3. Node 4 however did thicken in compacted soil and with ethylene exposure. However, root penetration for node 3 was difficult to assess as roots had shallow growth angles and hit the pot wall before interacting with the compacted layer, which reduced the number of replicates that could be sampled; and 3) Node 4 roots of IBM014 thickened when grown in soil, while they did not thicken with ethylene application.

Average cell size of genotypes grown in the hydroponics experiment were strongly correlated with cell size of those grown in soil (Figure 12, Table S3). The relationship between the soil and hydroponics experiments is stronger for node 3. Outer cortical cell area had lower R² values compared to those of middle and inner cortical cell area. Average cell size is slightly greater for node 4 roots together with greater standard deviations (Table S3). For node 3 genotype IBM014 had the greatest cell size in response to ethylene (Figure S8) and within the compacted layer (Figure 8). For node 4 roots of IBM086, the greatest cell size was attained in under growth in the compacted soil layer (Figure 7) and in ethylene treatments (Figure S8).

## Discussion

Literature suggests that cortical expansion of a root axis upon experiencing mechanical impedance is linked to ethylene, and genotypes that are responsive to ethylene would radially thicken (Moss *et al*., 1988; Sarquis *et al*., 1991). As root thickening relieves stress from the root tip (Bengough *et al*., 2006), it is often assumed that radial expansion will help roots to penetrate hard soil layers. In contrast to this expectation, in this study we observed that genotypes that showed less radial expansion upon encountering compacted soil were better able to cross a compacted layer and attained greater rooting depth than genotypes with greater radial expansion (Figure 6, S4). Furthermore, ethylene may be related to genetic variation in radial thickening since most genotypes showed similar anatomical responses to mechanical impedance conditions and exogenous ethylene application.

### Root thickening was driven by cortical cell size expansion rather than increased cell file number

Radial expansion upon encountering the compacted layer was mainly due to cortical expansion and, to a lesser extent, expansion of the stele (Figure 7) as the root cortical area is overall greater than the stele area. Depending on genotype and node, stele area increased or remained unchanged under impedance (Figure 7). Lupin roots that grew under impeded conditions maintained stele dimensions (Atwell, 1988; Hanbury and Atwell, 2005), while barley, maize, rice, pea and cotton roots showed increased stele diameters under impedance (Wilson *et al*., 1977; Iijima *et al*., 2007). Since the stele tissue is completely enclosed by the cortical tissue, radial expansion might be more difficult due to internal pressures between tissues restricting radial expansion. Alternatively, the cortex could simply be more plastic than the stele in its response to its local environment. Cortical tissues traits are responsive to other stresses (Chimungu *et al*., 2014a, 2014b; Saengwilai *et al*., 2014; Galindo-Castañeda *et al*., 2018), which illustrates the plasticity of this tissue. Huang *et al*. (1998) identified a cDNA clone (*pIIG1*) with higher expression in the cortical cells and protocambium of mechanically impeded maize roots illustrating that gene expression upon impedance can be localised in different root tissues. Functional consequences to drastic stele rearrangement could be important as xylem vessels might be affected as well as xylem vessel areas are correlated with stele area (Uga *et al*., 2008, 2009; Burton *et al*., 2015). For genotypes IBM014 and IBM086 we observed a significant increase in xylem vessel area in node 4 (Figure S9). How these changes affect water transport remains to be investigated.

Similarly to our results, Iijima *et al*. (2007) showed that the cortical thickness of maize increased more than that the stele diameter in response to mechanical impedance. Cortical changes due to impedance have been attributed to (1) increased cortical cell size (Atwell, 1988; Hanbury and Atwell, 2005; Veen, 1982) or (2) increase in both cell file number and cell size (Colombi *et al*., 2017; Croser *et al*., 1999; Iijima *et al*., 2007). These observations have used different plants, either exposed or not exposed to impedance, to obtain root axes for their observations. This would introduce additional uncertainty about cell file number changes. We have looked at anatomical changes along the axes of roots encountering impeding conditions, which has, to our knowledge not been done before. We observed that cortical thickening is due to cell diameter increases, while cell file number remained stable along the root axis (Figures 8, 9). Additionally, studies have documented species differences (Iijima *et al*., 2007; Colombi, 2016) rather than genotypic differences in response to mechanical impedance. Genotypic differences in anatomical response to mechanical impedance have only been studied in a few cases in wheat (Colombi, 2017, 2019) and maize (Chimungu *et al*., 2015; Vanhees *et al*., 2020).

The number of roots of different nodes within the same genotype crossing the compacted layer is not significantly different (Figure 3A, Table 3) although angle at which they encounter the layer might play a role (Figure 5). Node 3 and node 4 roots have more similar characteristics than nodes formed earlier and later, and earlier and later nodes may potentially differ in the proportion of roots able to overcome impedance conditions. This could be due to the innate difference in root cross sectional area, where thicker roots are predicted to experience less stress at the root tip and would experience smaller shear stresses over the root surface (Kirby and Bengough, 2002). Thicker roots are assumed to buckle less (Chimungu *et al*., 2015; Clark *et al*., 2003). We could however not test roots from other nodes in our current set-up due to pot-size and CT-scanner resolution limitations. Node 2 roots were hard to visualise and, because of their shallow growth angle, tended to encounter the pot wall before reaching the compacted layer. Roots younger than those of node 4 could not be evaluated because allowing plant growth beyond that stage would make evaluation difficult as columns become rootbound. Within roots from nodes 3 and 4, cross sectional area was not predictive for penetrability. Different wheat genotypes showed greater root penetration stress when root diameter increased under mechanical impedance (Colombi *et al*., 2017). Wheat plants have smaller diameter roots than maize. This difference in morphology could mean that wheat and maize could have different ways of dealing with impedance. Smaller diameter root axes may be able to explore the remaining porosity in a denser soil, while only lateral roots would be able to do so for maize (Cahn *et al*., 1989; Yamaguchi *et al*., 1990). The thicker roots of maize might have a competitive advantage when soil is unstructured as there will be fewer cracks or biopores to explore or when porosity is further reduced so that even thinner roots would experience mechanical stress. In these cases, thicker roots would be expected to experience less stress (Kirby and Bengough, 2002). Steeper root angles would allow roots to reach the layer within this pot system, but would also allow them to penetrate more easily as higher penetration rates were observed for steeper roots (Figure 4C). It could be that steeper roots are less likely to buckle when they encounter a harder soil layer, while roots that have a more shallow approach to the compacted layer might deflect more easily. However this remains to be investigated further as we were only able to sample a small range of root angles as roots that hit the pot wall, and thus were innately more shallow, could not be sampled. However, within our small range of root angles above the compacted layer we saw an effect of root angle on penetration rate, with those that were more steep having higher penetration rates (Figure 5).

Why roots thicken by cell size expansion rather than increasing their cell file number merits further study. Cortical cell expansion might be more energy efficient. Different wheat genotypes grown under impeding conditions all thickened and under greater impedance, genotypes with greater cortical cell diameters were more energy efficient (Colombi *et al*., 2019). A similar mechanism could form the basis for preferentially adjusting cell size instead of cell file number. Comparing similar root cortical areas composed of either greater number of cell files with smaller cells, or fewer cell files but with larger cell size, the latter may entail less metabolic cost to the roots, because of reduced cell wall construction, and the reduced metabolic costs of larger cells, which have been proposed to have reduced cytoplasm per unit tissue volume than smaller cells (Lynch, 2013; Chimungu *et al*., 2014a). Reduced metabolic costs assist with deeper rooting as the conserved resources can be used elsewhere in the plant including for greater soil exploration (Lynch and Wojciechowski, 2015; Lynch, 2015). In addition, a change in cell size may be easier and quicker to achieve than a cell file number change which would entail meristematic reorganization.

Cortical cell size varied across the cortex (Figure 8) and outer cell layers expanded to a proportionally greater extent in the compacted layer (Table 6). For wheat and maize, greater outer cortical cell expansion has been reported in response to mechanical impedance (Wilson *et al*., 1977; Veen, 1982). Why the different regions expand differentially remains unclear. Expansion of outer cortical layers may be less limited as they experience less internal pressure from surrounding cells (Bengough *et al*., 2006; Veen, 1982). Outer cortical cells remained smaller than middle cortical cells (Figure 8) and it has been suggested that several layers of smaller cells in the outer region of the cortex provide mechanical stability (Chimungu *et al*., 2015; Striker *et al*., 2007). The inner and middle cortex of maize primary roots was observed to be more sensitive to exogenous ethylene than the outer cortex, with greater radial expansion at the expense of elongation (Baluška *et al*., 1993). In our experiment, ethylene treatment caused similar cell size expansion across the cortical regions though this was not the case for roots grown in compacted soil (Table 6). Our results could be different from those of Baluška *et al*. (1993) because primary and nodal roots behave differently or because our plants were exposed to continuous ethylene treatment throughout development as opposed to 24h in the other study.

### Root thickening did not improve root penetration through a compacted soil layer

Ethylene appears to be involved in the radial thickening response, since the genetic variation in ethylene-induced thickening was correlated with the genetic variation in impedance-induced thickening (Figure 11). Impeded roots produce more ethylene than non-impeded controls (Moss *et al*., 1988; Sarquis *et al*., 1991; He *et al*., 1996). Root cross sectional area measured on roots above the compacted layer (experiment 1) and those under control conditions and 1-MCP treatment (experiment 2) were comparable (Figure 11). 1-MCP should block ethylene perception by roots, and exhibited significant effects on root morphology (Figures S6). It can therefore be assumed that thickness of roots growing through less impeding soil (before and after the compacted layer) were not significantly influenced by ethylene. If the ability to cross the compacted layer was solely due to enlarged root diameter, all roots would need to radially expand to a certain diameter regardless of genotype or node. This was not the case, for example node 4 roots of IBM146 had the smallest root cross sectional area within the compacted layer (Figure S4), while having the greatest penetration rate (Figure 3) and the steepest root angle (Figure 4). Furthermore we observed swollen root tips on those roots that buckled when encountering the compacted layer, which further illustrates that thickening does not always enable penetration of the layer.

Ethylene regulates root extension and lateral root density (Figure S6; Moss *et al*., 1988; Sarquis *et al*., 1991; Borch *et al*., 1999). Root thickening is associated with reduced elongation rates (Bengough and Mullins, 1991; Croser *et al*., 2000) through the reduction of cell length and cell flux out of the meristem (Croser *et al*., 1999). Ethylene itself reduces the number of meristematic cells, which reduces meristem length (Barlow, 1976). Ethylene also reduces cell elongation and increases radial expansion, resulting in less root elongation (Sarquis *et al*., 1991) and promotes root hair cell elongation (Pitts *et al*., 2001), which could stabilise the root and help penetration (Haling *et al*., 2013; Bengough *et al*., 2016). Our study suggests that roots that are ethylene insensitive can maintain root elongation under impeded conditions, enabling them to attain greater rooting depth and potentially allowing better access to water and nutrients in deep soil strata. However, positive effects have also been attributed to root thickening. For instance, thickening reduces the stress on the root tip (Kirby and Bengough, 2002) and thicker roots buckle less (Clark *et al*., 2008; Whiteley *et al*., 1982). Thickening of roots might be beneficial on small scales or for localised impeded conditions. In order for roots to penetrate harder soil clods/aggregates or to penetrate through a biopore wall, usually only a small distance of impedance needs to be overcome. However, the effect of thickening and reduced elongation rate clearly leads to reduced root length and soil exploration by the affected root axis. We propose that the negative effects of ethylene will increasingly overrule the positive with increasingly thick layers of compacted soil.

Moss *et al*. (1988) found that application of ethylene reduced primary root length further the longer it was applied. Under prolonged impeded conditions, ethylene, as a stress signal, could potentially inform the plant to alter its growth by compensatory root growth mechanisms. The compacted layer in this research was designed to mimic the spatial abruptness of a plough pan, which could induce different anatomical responses than when a root axis has been experiencing impedance for a longer time. How prolonged exposure to impedance, for instance when growing through compacted soil instead of a hardpan, changes root anatomy and root architecture within a whole root system and how this differs from the current experimental system remains to be investigated. We observed that anatomical phenotypes recovered once the root had passed the compacted layer. Similarly, root elongation rates of barley were restored after 3 days when transferred from impeded conditions in ballotini to unimpeded growth in solution (Goss and Russell, 1977) and pea roots experienced reduced elongation rates for 48 hours after transferring to hydroponics after which the former elongation rate was restored (Croser *et al*., 2000). Assuming that under unimpeded conditions these roots can elongate more than 1 cm per day, we saw that the residual effect of impedance in soil was less pronounced than in other studies. Ethylene production rates can rapidly increase and decrease upon application of mechanical impedance (Sarquis *et al*., 1991). The concentration of ethylene that roots are exposed to also plays a role as higher ethylene concentrations induce longer recovery time (Whalen and Feldman (1988) cited by Sarquis *et al*. (1991)). Under our experimental conditions, the change in soil penetration resistance was 0.35 MPa, less than in most other studies. It would therefore be reasonable to assume that a short-term ethylene signal was present, after which roots quickly return to their original radial dimensions. It is also likely that roots will have experienced a range of physical stresses within the compacted layer, as the soil dried and then was re-wet, following watering. This may have significantly increased the degree of mechanical impedance when the soil was drier, and perhaps even permitted transient hypoxia following rewatering.

We suggest that ethylene functions as a stop signal for root growth when axial roots become impeded (Pandey et al. 2021). When larger volumes of impeded soil cause a prolonged production of ethylene, this will signal axial root growth to stop. Upon this signal, root growth in the lesser impeded areas, or adjustments to above ground plant growth might become upregulated.

## Conclusions

Root thickening within a compacted layer varied with genotype. Previous studies have not considered anatomical changes along individual root axes in response to impeding soil conditions. We found no significant changes to the cell file number along a single root axis of maize when this axis grew through denser soil. Instead, thickening of the cortex was caused by cell radial expansion. Exogenous ethylene and mechanical impedance caused similar patterns of expansion in cortical cells. Root thickening negatively correlated with the ability of the different genotypes to penetrate through a compacted soil layer and grow past the compacted layer. Genotypes that did not thicken when encountering the compacted layer or under application of exogenous ethylene had the highest penetration percentages and were able to grow deeper past the compacted layer. This was node and genotype dependent. As root thickening is associated with reduced elongation rates, we suggest that prolonged exposure to ethylene slows and may ultimately stop axial root growth. This implies that ethylene will stop further root exploration when roots experience impedance and that roots with less ethylene responsiveness could be better at overcoming impedance in many situations.

## Acknowledgements

The authors thank Brian Atkinson and Craig Sturrock for assistance with the X-ray CT-study. This research was supported by the University of Nottingham, the Pennsylvania State University and the James Hutton Institute. The James Hutton Institute receives funding from the Rural & Environment Science & Analytical Services Division of the Scottish Government.

## Supplementary data

### Figures

**Figure S1.**
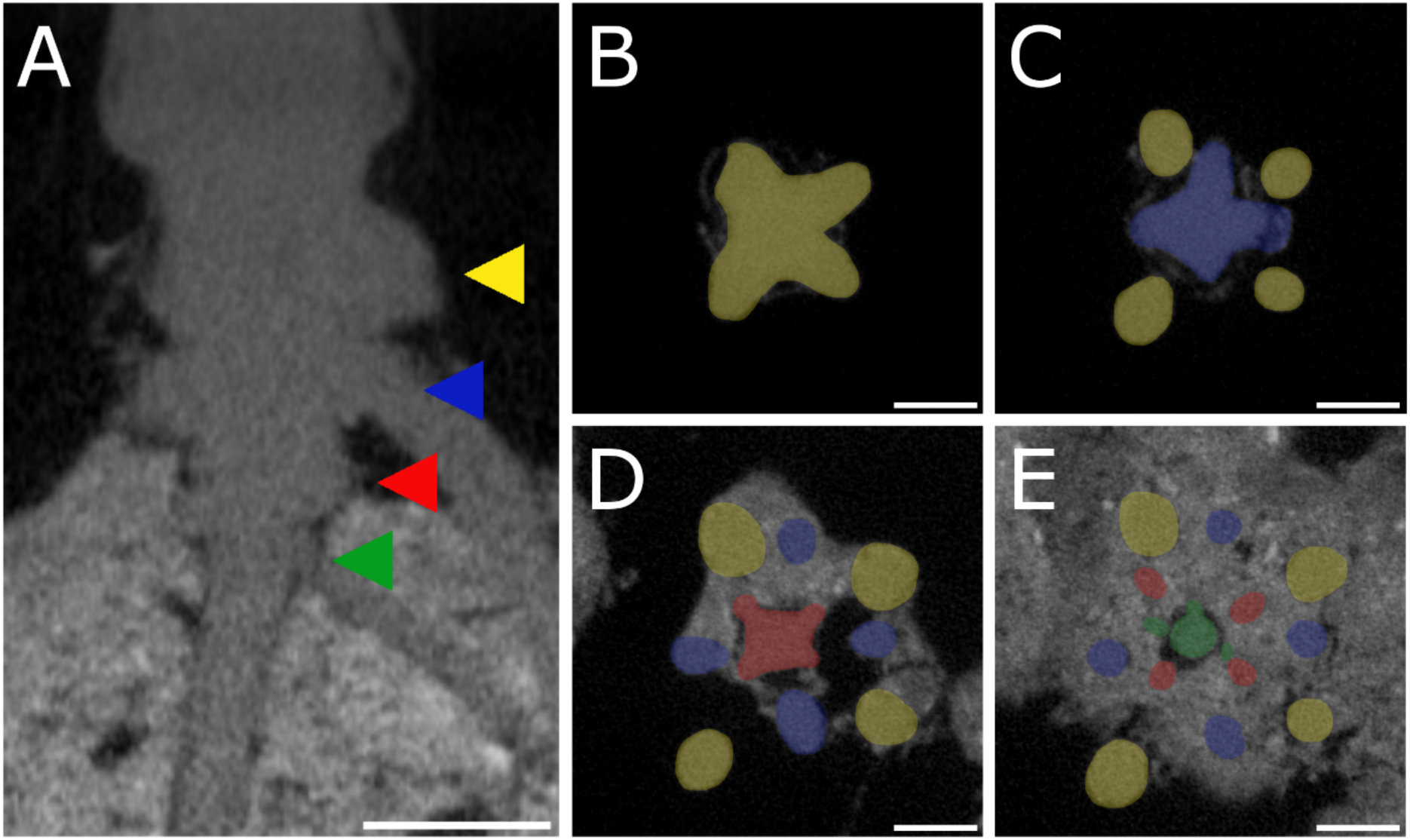
Node identification on 2 dimensional planes during image processing of X-ray CT scans. (A) shows a xy-projection at the root base. (B-E) show different yz-projections moving from the top of the column down. Different nodes are indicated by the different colours (green – node 1, red – node 2, blue – node 3, yellow – node 4). Scale bars are set at 1 cm.

**Figure S2.**
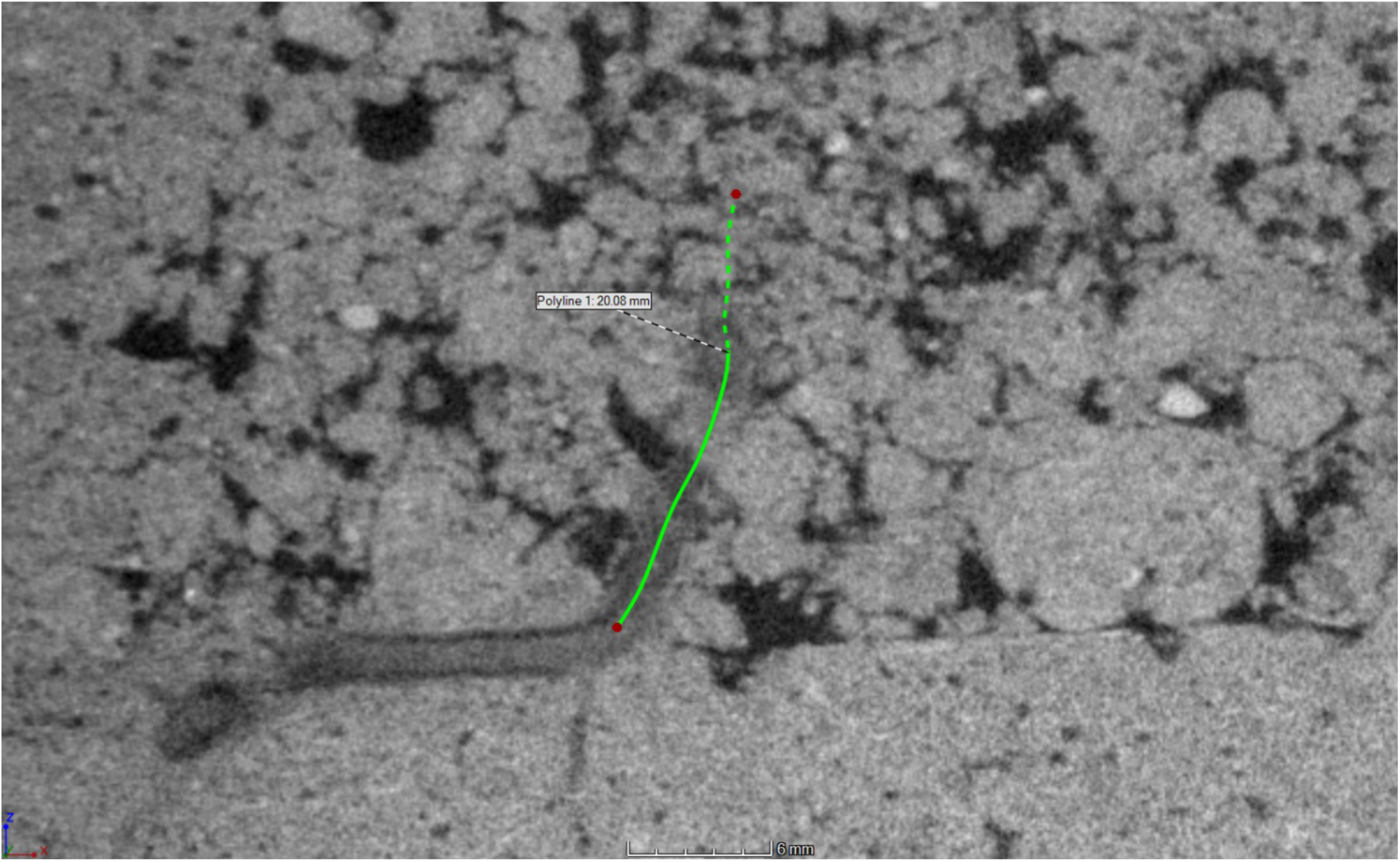
Example of a polylined root segment of approximately 2 cm of a deflecting nodal root upon the layer. The (dotted) green line represents the projection of the polyline onto the xy-plane.

**Figure S3.**
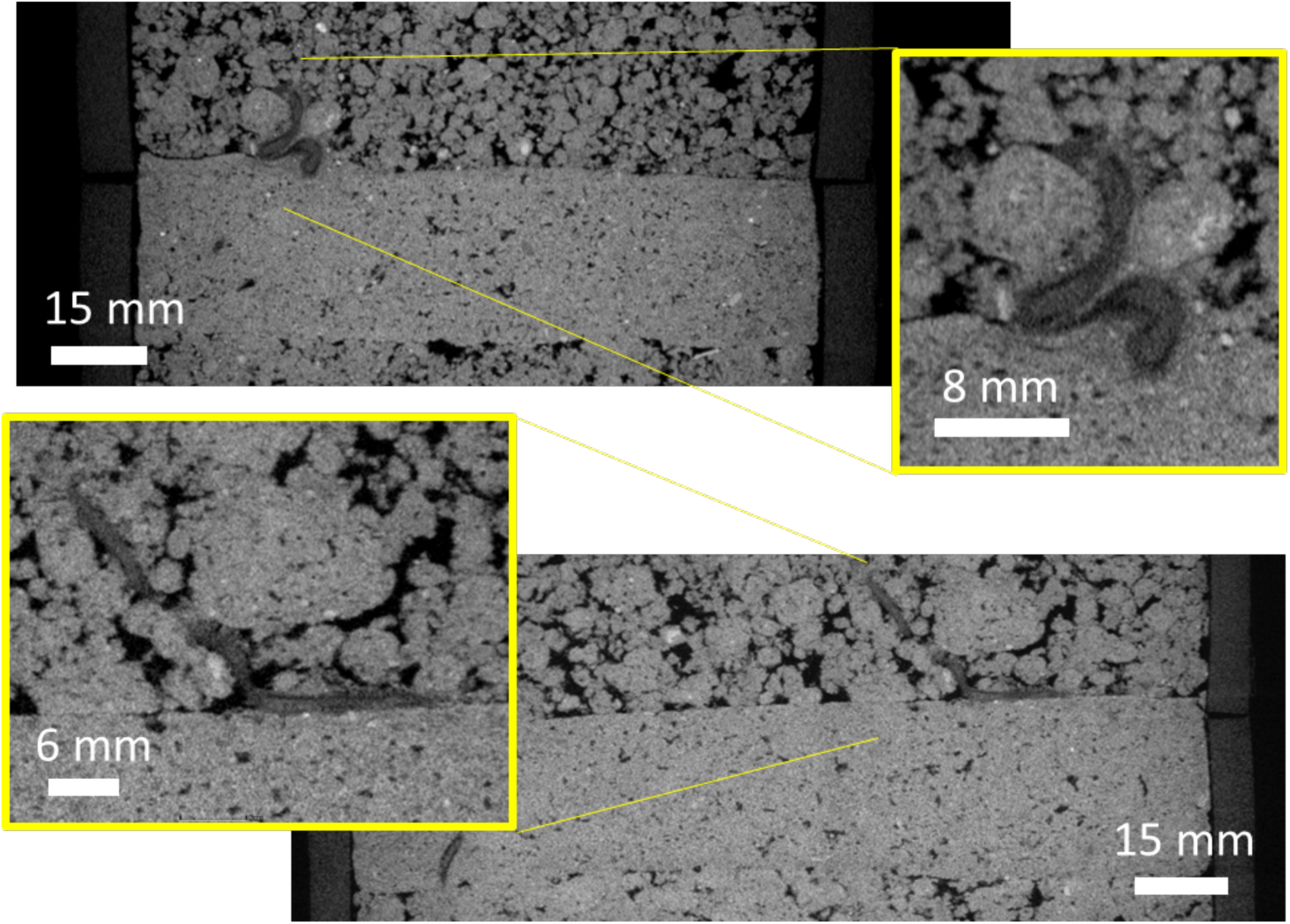
Nodal roots of maize can buckle (top panel) or deflect (bottom panel) when encountering a dense layer.

**Figure S4.**
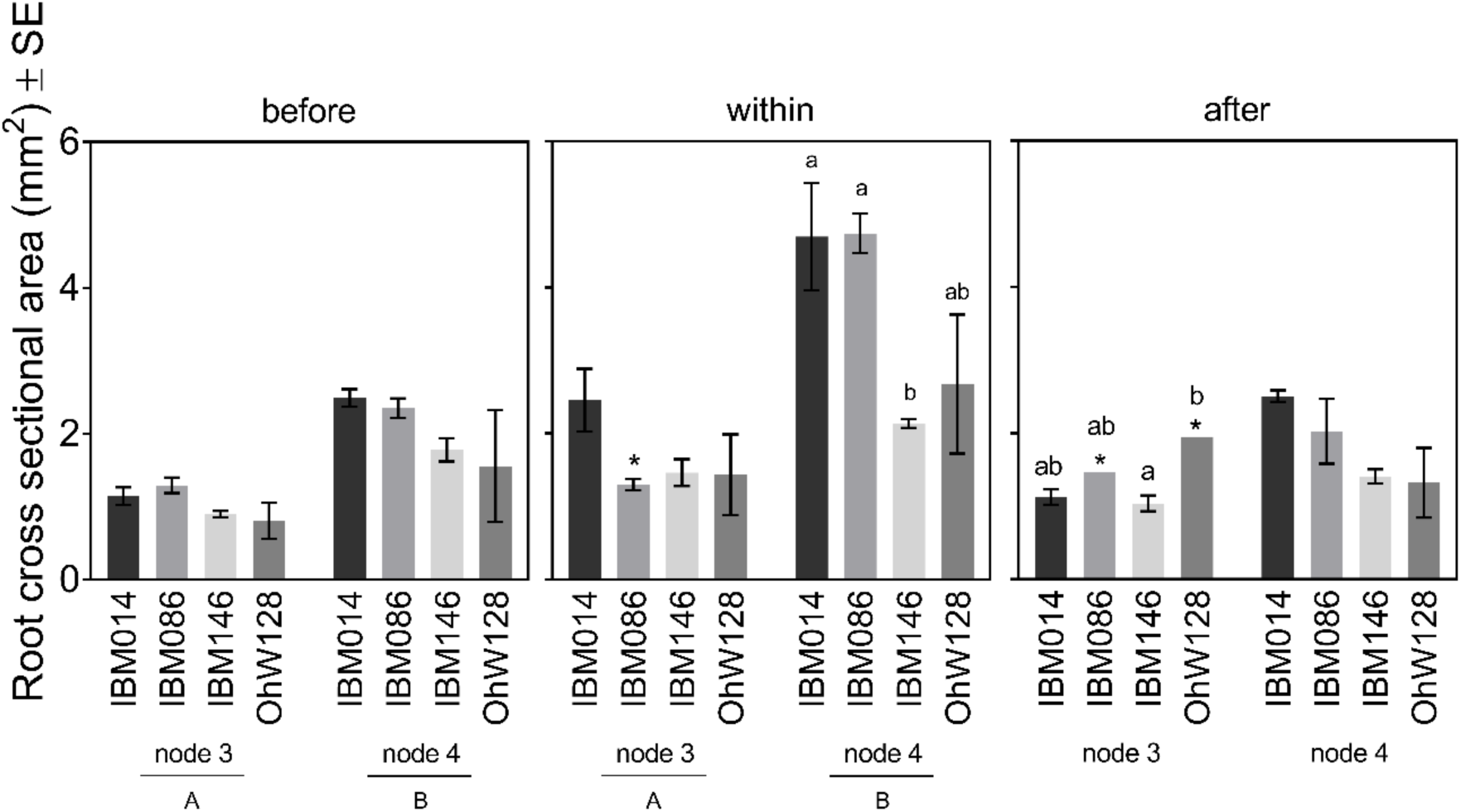
Root cross sectional area for both nodes and four genotypes before, within and after the compacted layer. Differences between nodes (capital letters, P ≤ 0.001) and between genotypes within respective nodes (lower case letters, P ≤ 0.05) were calculated by Tukey comparisons. Genotypes indicated by * had a limited amount of sections due to limited amount of roots able to cross the compacted layer. Where no letters are shown, no significant differences were found between nodes or genotypes within nodes.

**Figure S5.**
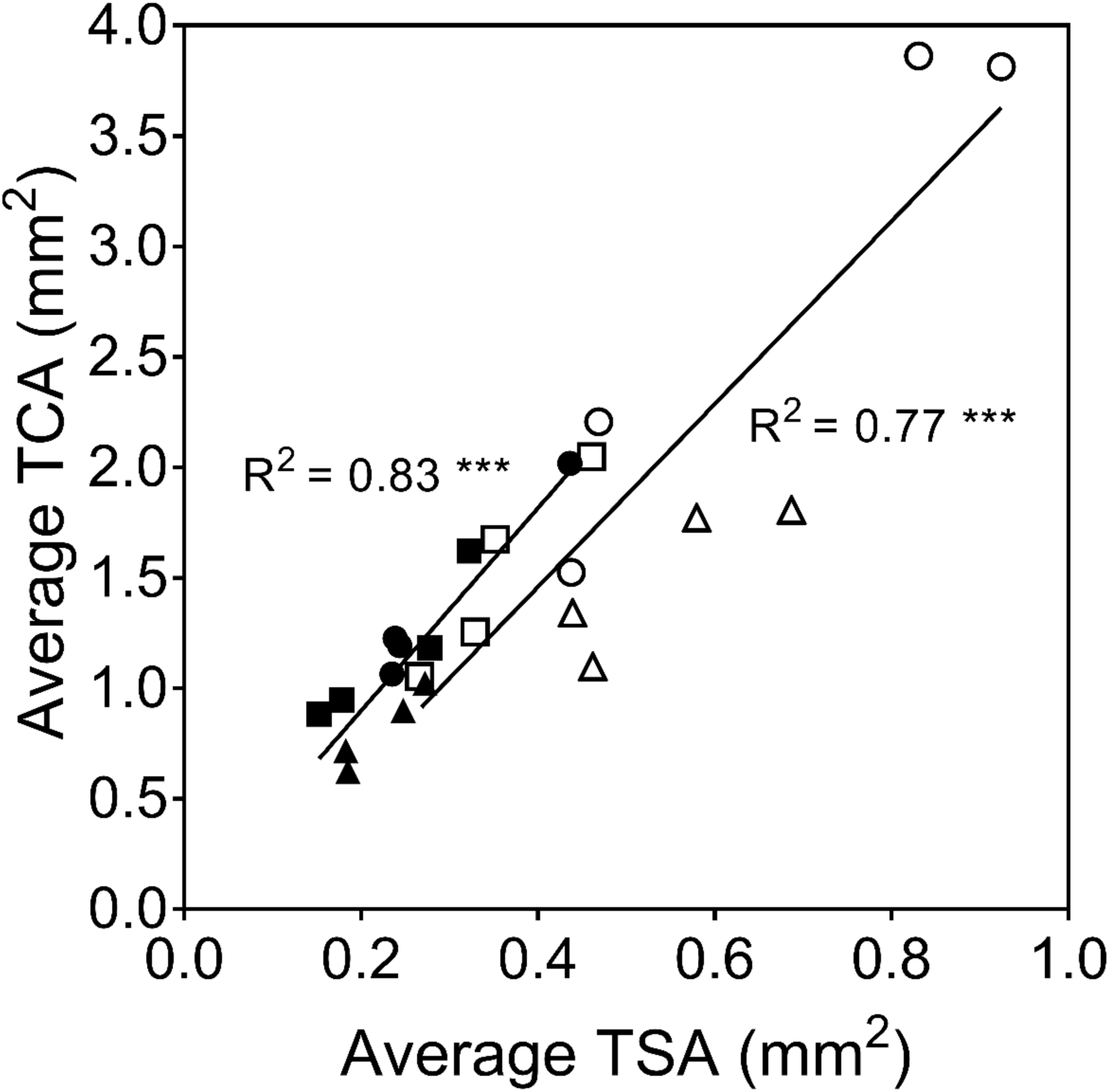
Correlation between stele area and cortical area before (triangles), within (circles) and after (squares) the compacted layer for node 3 (black symbols) and node 4 (white symbols). Level of significance at p ≤ 0.001.

**Figure S6.**
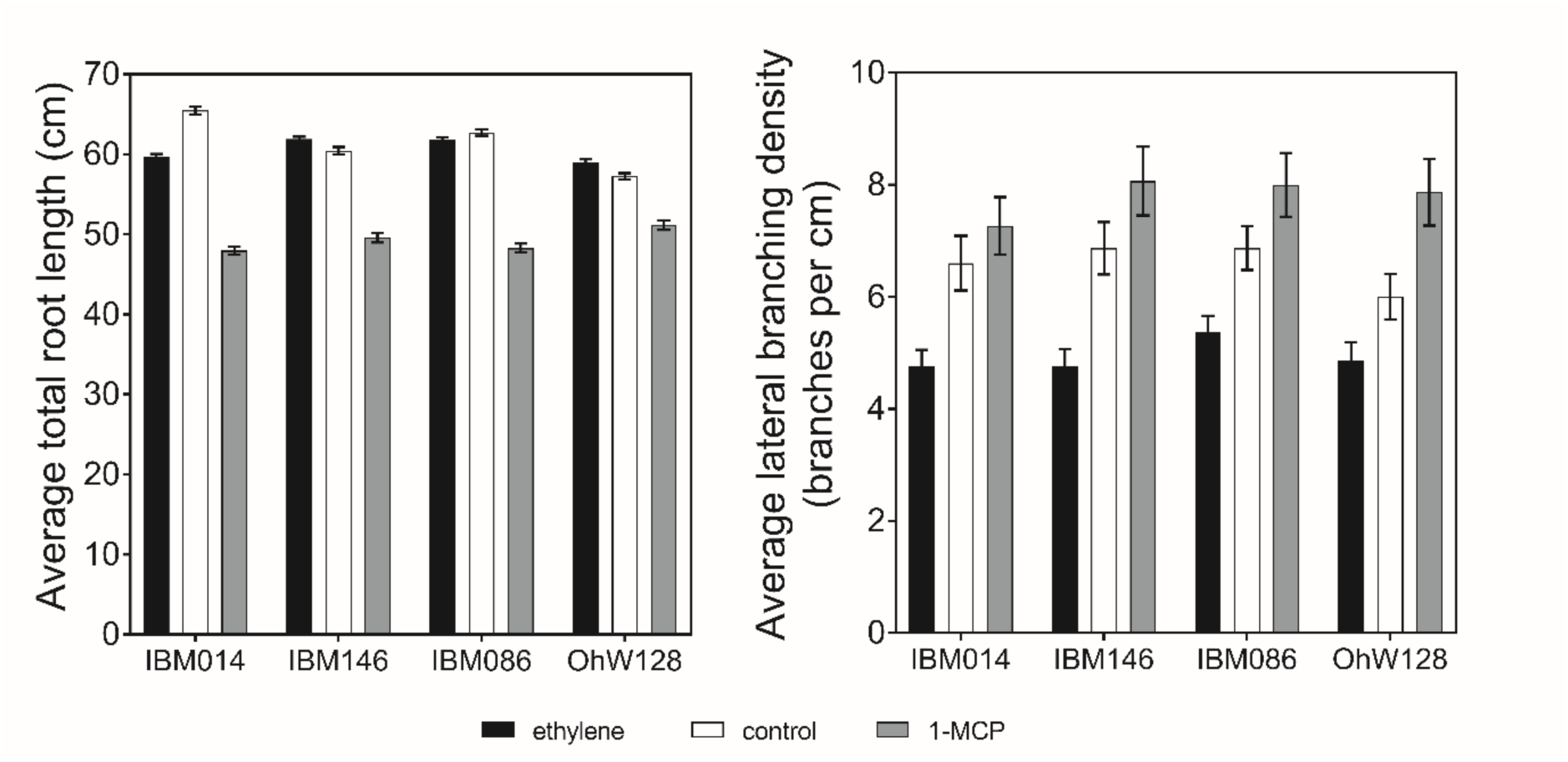
Average total root length (cm) ± SE and average lateral branching density (branches per cm) ± SE for the four different genotypes tested under ethylene treatment, 1-MCP treatment and control.

**Figure S7.**
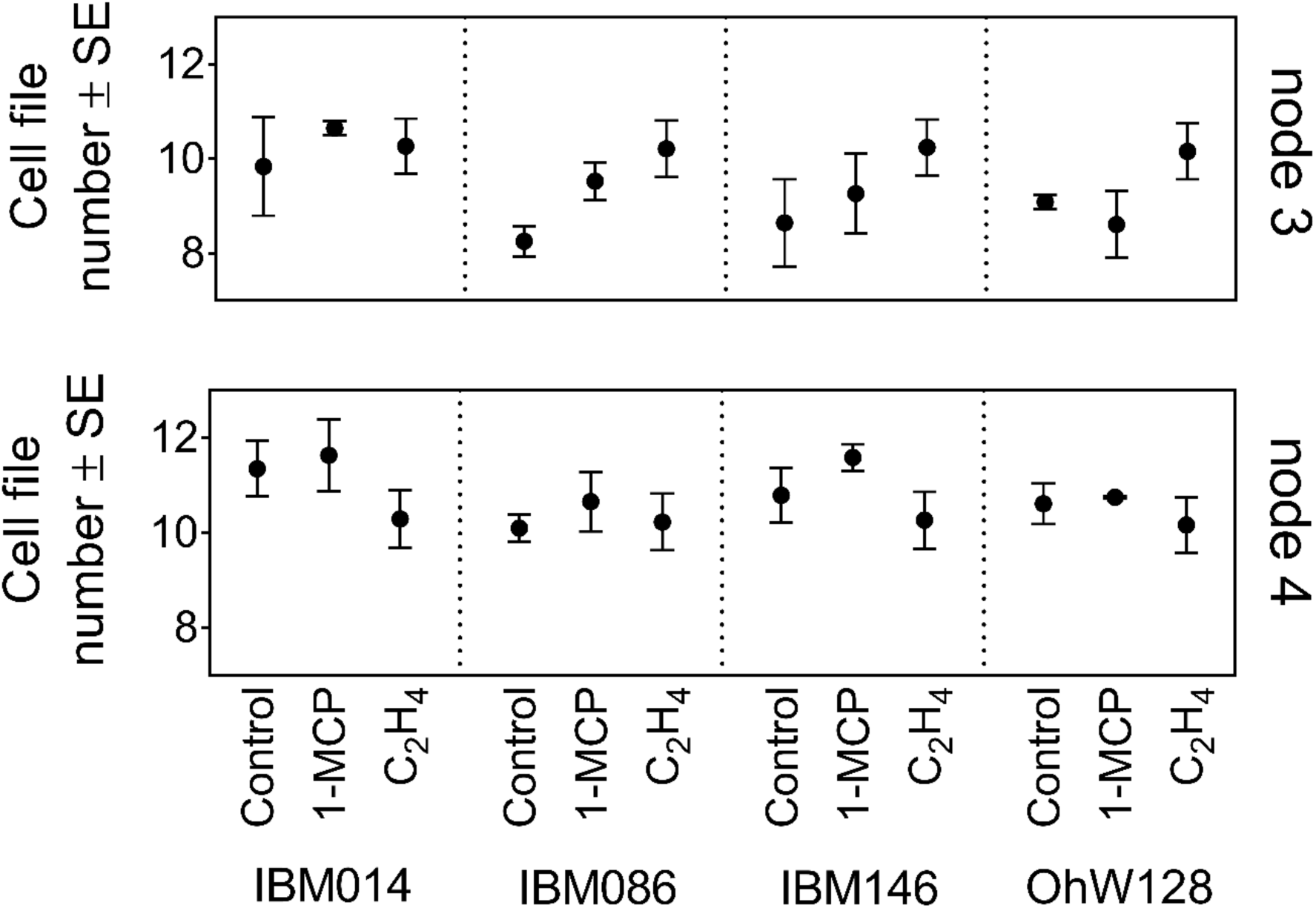
Average cell file number ± SE for different nodes and genotypes under ethylene treatment. No significant differences were found between treatments within each genotype-node combination. For some observations the standard error was so small it could not be visualised.

**Figure S8.**
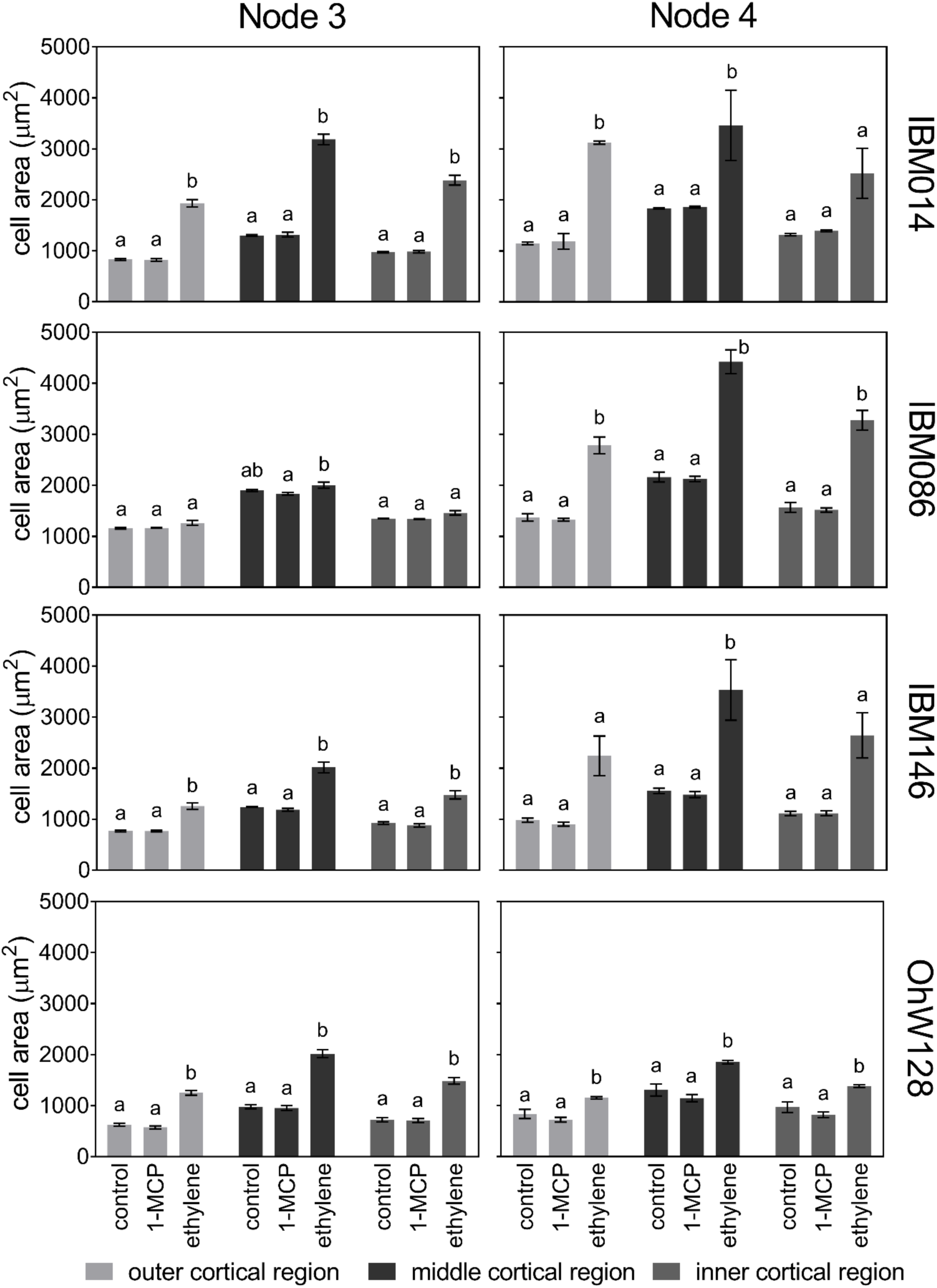
Average cortical cell size (µm²) ± SE for different cortical cell positions within root cross sections when either 1-MCP or ethylene was applied to the root system versus a control. Differences among treatments were calculated by Tukey comparisons within node - genotype combinations (P ≤ 0.05).

**Figure S9.**
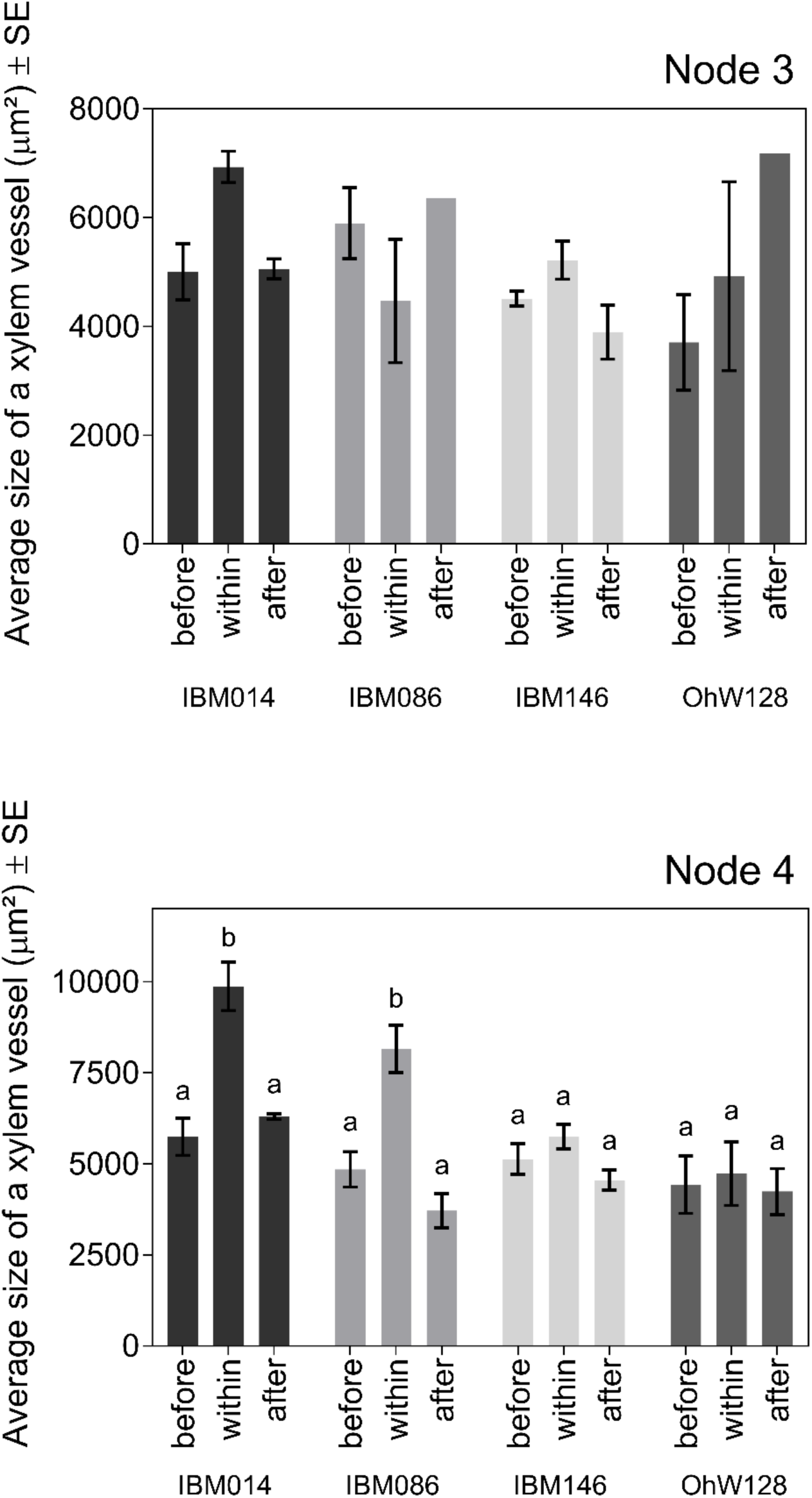
Average xylem vessel areas for each genotype and each node. No significant differences were found for node 3 for xylem vessel area before, within and after the compacted layer. For node 4 there were significant differences identified with Tukey comparisons (P ≤ 0.001).

### Tables

**Table S1.**
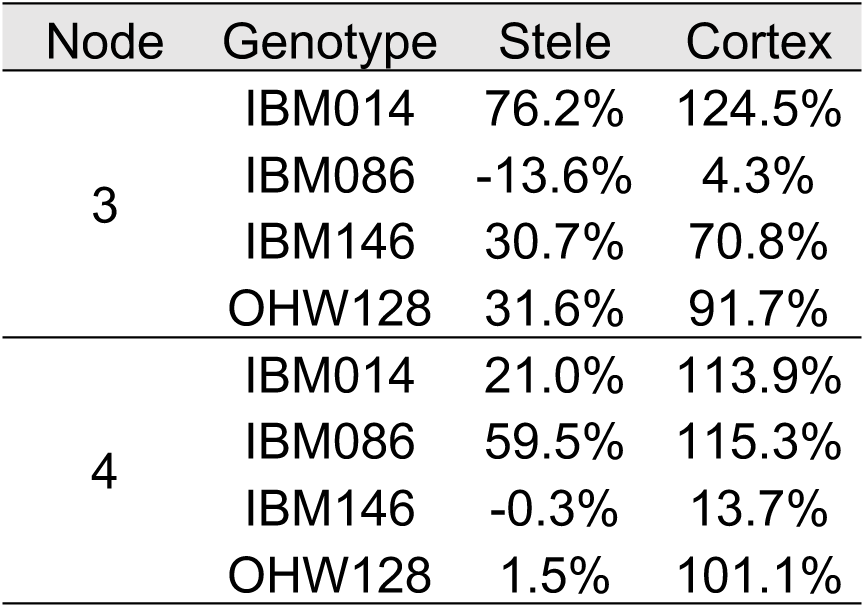
Relative increase or decrease in cortical or stele area when roots grow from above the layer into the compacted layer.

**Table S2.**
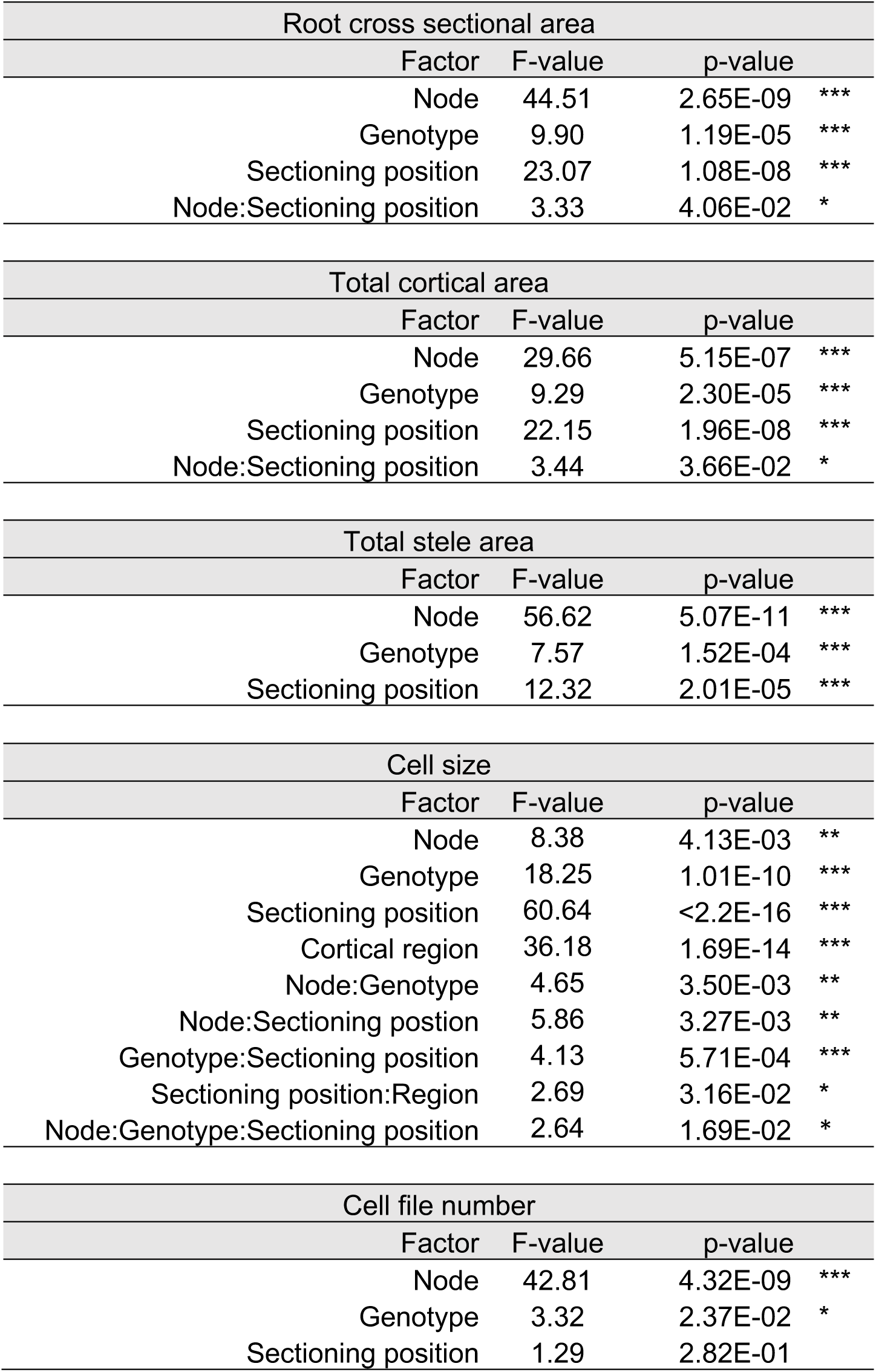
ANOVA results for anatomical traits. Each table shows all the main effect results regardless of significance, interaction terms were discarded if proven insignificant.

**Table S3.** Average cortical cell area per cortical region per node in soil or ethylene.

|  | node 3 |  |  |  | node 4 |  |  |  |
| --- | --- | --- | --- | --- | --- | --- | --- | --- |
|  | soil |  | ethylene |  | soil |  | ethylene |  |
| Outer | 1139 | ± 434 | 1136 | ± 408 | 1392 | ± 1008 | 1585 | ± 714 |
| Middle | 1930 | ± 846 | 1830 | ± 686 | 2579 | ± 1506 | 2516 | ± 1133 |
| Inner | 1286 | ± 429 | 1347 | ± 512 | 1663 | ± 830 | 1849 | ± 846 |

